# *Rubus armeniacus* genome sequence reveals the secrets of blackberry anthocyanin biosynthesis

**DOI:** 10.64898/2026.05.05.723051

**Authors:** Katharina Wolff, Melina Sophie Nowak, Corinna Thoben, Till Beuerle, Boas Pucker

## Abstract

Here, we present a comprehensive multiomics analysis of anthocyanin biosynthesis in *Rubus armeniacus*, known for its dark fruits. A phased genome sequence of the tetraploid blackberry was generated, achieving an N50 of 34 Mb with an assembly size of 1.2 Gbp based on Oxford Nanopore Technology sequencing (ONT). The BUSCO score for the total assembly shows a high completeness of 99.1%. The assembly was separated into 4 pseudohaplophases, with the pseudohaplophase A representing the *R. armeniacus* genome in 7 chromosome scale contigs, with an N50 of 46 Mbp and 98.8% conserved BUSCO genes. A total of 118,183 protein coding genes were annotated within the genome assembly and all relevant genes encoding enzymes and transcriptional regulators of the anthocyanin biosynthesis pathway were identified within each pseudohaplophase. To further understand the underlying cause of dark pigmentation, the gene expression was analysed during different stages of berry development revealing a strong induction of anthocyanin biosynthesis genes including the anthocyanin activating subgroup 6 MYB transcriptions during the berry ripening process. Further, a quantification of cyanidin-3-O-glucoside in methanolic berry extract, utilizing a UHPLC-HRAM-MS analysis, revealed an approximately 500-fold increase of cyanidin-3-O-glucoside from green to black fruit, indicating that dark pigmentation in *R. armeniacus* results from high anthocyanin accumulation.

**Significance statement:** This study provides a multiomics analysis of the dark pigmentation of *Rubus armeniacus,* including a high quality phased assembly and an in-depth analysis of the anthocyanin biosynthesis pathway. A transcriptional and metabolomic analysis revealed that dark berry pigmentation is caused by a high accumulation of cyanidin-3-O-glucoside during fruit ripening.

## Introduction

The genus *Rubus* includes several crops with great economic interest such as the red raspberry (*Rubus idaeus*) and black raspberry (*Rubus occidentalis*) both belonging to the subgenus *Idaeobatus.* For this subgenus several genomic resources for different species are available and have been studied intensely (Brůna et al., 2023). However, the availability of full genome sequences for the subgenus *Rubus*, with dark coloured fruits is still underrepresented, despite including economically interesting crops such as the blackberry. One of the few species of the *Rubus* subgenus which has been sequenced is *Rubus argutus* “Hillquist”, a diploid blackberry cultivar, which is commonly used in breeding and crop production, mostly known for its trait of primocane-fruiting (Brůna et al., 2023). Within the *Rubus* subgenus, ploidy can vary from diploid up to dodecaploid (12x) with most commercially used cultivars being tetraploid. Recently, the first haplophase resolved genome assembly of the thornless tetraploid blackberry *Rubus* L. subgenus *Rubus Watson* has been published (Paudel et al., 2025). Based on these genetic resources, candidate genes for several desirable breeding traits in blackberry, such as annual flowering, thornlessness or disease resistance genes were identified (Brůna et al., 2023; Paudel et al., 2025). For example, the WUSCHEL-LIKE HOMEOBOX transcription factor, WOX1, was identified as a key regulator in the development of prickles in blackberry and raspberry species (Aubin et al., 2026). Sequencing of wild species such as *R. armeniacus* offers an opportunity to broaden the genetic base, facilitate the discovery of novel traits, and enhance understanding of the evolutionary history and diversity within the *Rubus* genus.

Beyond genetic and breeding studies, the biochemical properties of blackberry fruit—particularly the accumulation of anthocyanins—have garnered increasing attention due to their contribution to fruit quality and anthocyanins association with human health benefits, based on their antioxidant and anti-inflammatory properties (Ma et al., 2021). In plants, they are involved in protection mechanisms against abiotic and biotic stresses (Grünig et al., 2025; Naing et al., 2017; Saad et al., 2021). Additionally, anthocyanins are widely distributed among angiosperms and can protect plants against high light irradiation (Landi et al., 2015; Tattini et al., 2014) and serve as pigments in leaves, flowers, and fruits (Winkel-Shirley, 2001). Due to their colorful properties, anthocyanins are involved in attracting animals for pollination and seed dispersal, thereby contributing to reproduction processes in plants (Grünig et al., 2025). Anthocyanins are responsible for a variety of vibrant colours such as pink, red, blue and purple in various plant tissues and can even cause darker pigmentation. Several studies have been undertaken to understand the underlying mechanism of dark pigmentations in various plant species such as *Aronia melanocarpa* (Mahoney et al., 2022) and *Vaccinium myrtillus L.* (Lätti et al., 2008; Plunkett et al., 2018) and revealed a high concentration of anthocyanins in the respective tissue. The pigmentation of the *Rubus* genus has been previously reported to be based on anthocyanins with the main anthocyanin present being cyanidin-3-O-glucoside (Toshima et al., 2021). However, there appear to be species-specific differences on the identity of these anthocyanins and their importance and contribution to the dark colouration. While the dark pigmentation within petals or foliage is rare, dark pigmentation in berries can be observed at a higher rate, but the mechanism behind this pigmentation is not yet fully understood (Blando et al., 2019; Weatherall and Lee, 1991; Wolff and Pucker, 2025). Dark pigmentation can serve several biological functions for the plant such as the attraction of pollinators, in the form of petal spots and in the context of mimicry or sexual deception (Ellis et al., 2014; Thomas et al., 2009). Further studies have shown that high levels of anthocyanins can serve as a form of protection of the photosynthesis apparatus by the absorbance of UV-A and UV B rays (Gould, 2004; Landi et al., 2015).

Blackberries exhibit a distinctive dark pigmentation, which has been attributed to high concentration of anthocyanins (Bowen-Forbes et al., 2010), making them an ideal model species to study the accumulation and regulatory mechanisms of the anthocyanin biosynthesis pathway. In general, anthocyanins belong to a versatile group of specialized plant metabolites: the flavonoids.

The synthesis of anthocyanins begins with the general phenylpropanoid pathway comprised of three enzymes, namely: phenylalanine ammonium lyase (PAL), cinnamate 4-hydroxylase (C4H), and 4-coumarate:CoA ligase (4CL). These enzymes facilitate the conversion of phenylalanine to p-coumaroyl-CoA, the first substrate of the flavonoid biosynthesis. Naringenin-chalcone synthase (CHS), chalcone isomerase (CHI) and flavone 3-hydroxylase (F3H) catalyse the reactions to dihydrokaempferol. From dihydrokaempferol, the anthocyanin pathway splits into three parallel branches, based on the addition of hydroxyl groups to the B ring of the molecule: flavonoid 3’-hydroxylase (F3’H) and flavonoid 3’,5’-hydroxylase (F3’5’H), leading to cyanidin or delphinidin derivatives, respectively, while pelargonidin derivatives are utilizing dihydrokaempferol as the substrate (Seitz et al., 2007). The last four enzymes are conserved between the different anthocyanin classes and consist of the dihydroflavonol 4-reductase (DFR), anthocyanidin synthase (ANS), and anthocyanin-related glutathione S-transferase (arGST) to produce anthocyanidins which are converted into anthocyanins through the UDP-dependent anthocyanidin-3-O-glucosyltransferase (F3GT) based on the addition of a sugar moiety (Eichenberger et al., 2023; Grotewold, 2006; Winkel-Shirley, 2001). Additionally, the anthocyanin regulation has been largely attributed to MYB transcription factors of the Subgroup 6 and certain bHLH, particularly TT8, in combination with the WD40 protein TTG1, forming the MBW-complex (Gonzalez et al., 2008; Marin-Recinos and Pucker, 2024).

This study presents a fully phased genome assembly for the wild blackberry species *Rubus armeniacus* and the annotation of the anthocyanin biosynthesis genes, as well as a detailed investigation of the expression of anthocyanin biosynthesis genes and transcription factors in different ripening stages of berries revealing the genetic basis of color change during berry development.

## Results

### Assembly results

For the generation of the genome assembly, a total of 806,016 reads of ONT R9.4.1 and 2,190,500 reads of ONT R10.4.1 sequencing data were generated, respectively. The dataset comprises a total of 68.47 Gbp with an N50 value of 32.7 kbp. The genome size of *R. armeniacus* is estimated to be 300 Mbp, based on the genome assembly of its close relative *R. argutus* (Brůna et al., 2023). Given the amount of data generated the expected average coverage depth of the genome is 228.

Three different assemblers were utilized to generate the genome assembly, namely NextDenovo2, Hifiasm, and Shasta (Table 1). The following assemblies were generated based on the combined R9.4.1 and R10.4.1 reads corrected by HERRO.

**Table 1:** Assembly statistics for the assemblies of *Rubus armeniacus* based on NextDenovo2 (RAN), Hifiasm (RAH), and Shasta (RAS), including the total number of contigs, minimum and maximum contig length, N50 as well as BUSCO score (C: complete, S: single copy, D: Duplicated, F: Fragmented, M: Missing BUSCO genes) based on the rosaceae_odb12 dataset.

| assembler | NextDenovo2 (RAN) | Hifiasm (RAH) | Shasta (RAS) |
| --- | --- | --- | --- |
| assembly size [bp] | 1,196,153,256 | 986,225,330 | 1,253,341,178 |
| total contigs | 382 | 172 | 65 |
| min. contig length [bp] | 50,729 | 50,252 | 50,726 |
| max. contig length [bp] | 31,386,151 | 60,154,785 | 60,730,349 |
| N50 [bp] | 12,801,236 | 39,474,006 | 35,485,994 |
| BUSCO score | C:99.5%<br>[S:8.3%,D:91.2%]<br>,F:0.0%,M:0.4% | C:99.1%<br>[S:0.4%,D:98.7%],<br>F:0.3%,M:0.6% | C:99.1%<br>[S:0.2%,D:98.9%],<br>F:0.3%,M:0.6% |

Both the assembly from NextDenovo2 (RAN) and the Shasta assembly (RAS) produced assemblies of similar sizes of 1.20 Gbp and 1.25 Gbp, respectively, while the assembly with Hifiasm (RAH) resulted in a decreased assembly size of 0.99 Gbp.

The assembly RAN consists of 382 contigs with an N50 value of 12.80 Mbp. BUSCO was applied to assess the completeness of the assembly. In total 99.5% of the 10071 BUSCOs included in the rosaceae_odb12 dataset were present in the RAN assembly. Only 8.3% of the found BUSCOs were present as single copies, while the remaining 91.2% were duplicated.

The assembled genome sequence in the RAH assembly is represented by 175 contigs, with an N50 value of 38.47 Mbp and a BUSCO completeness of 99.1 % with a high BUSCO gene duplication rate of 99.1 %.

For RAS, a total of 65 contigs were assembled with an N50 value of 35.49 Mbp. The assessment based on BUSCO shows a completeness of 99.1% with 0.2% of BUSCO genes being present in the assembly as a single copy while 98.9% were represented as duplicates. Due to the reduced amount of contigs as well as the largest genome assembly size of RAS, this assembly was selected as the representative genome sequences and was annotated and further analysed.

### Ploidy analysis and separation of haplophases

The RAS genome assembly includes a total of about 1.2 Gbp, which is 4 times higher than the expected haploid genome size of 300 Mbp, as well as the analysis of the assembly revealed a high duplication of BUSCO score indicating a phased or partially phased assembly. In order to determine the ploidy of the assembly the amount of duplication of each BUSCO gene was analysed (Figure 2A), revealing that out of 1614 BUSCO genes 1440 are represented in the assembly in 4 copies. This indicates that RAS represents the tetraploid genome sequence of *R. armeniacus*.

**Figure 1:**
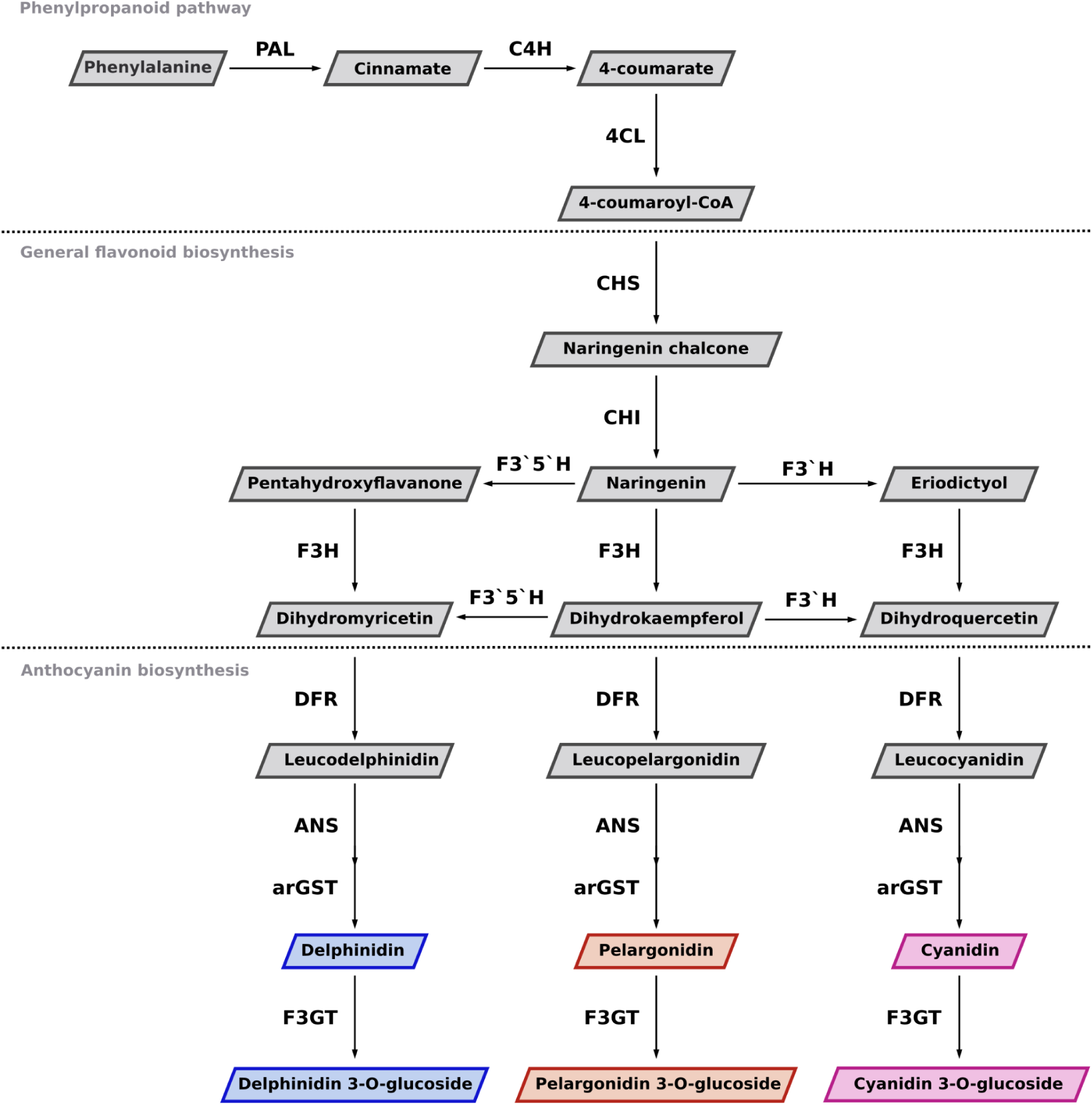
Schematic illustration of the phenylpropanoid (above dashed line), flavonoid biosynthesis (middle) and anthocyanin biosynthesis (below second dashed line) pathway. PAL (Phenylalanine ammonia lyase), C4H (Cinnamate 4-hydroxylase), 4CL (4-coumarate-CoA ligase), CHS (Naringenin-chalcone synthase), CHI (Chalcone isomerase), F3H (Flavanone 3-hydroxylase), F3’H (Flavonoid 3’-hydroxylase), F3’5’H (Flavonoid 3’,5’-hydroxylase), DFR (Dihydroflavonol 4-reductase), ANS (Anthocyanidin synthase), arGST (anthocyanin-related Glutathione S-transferase), F3GT (UDP-dependent anthocyanidin-3-O-glucosyltransferase).

**Figure 2:**
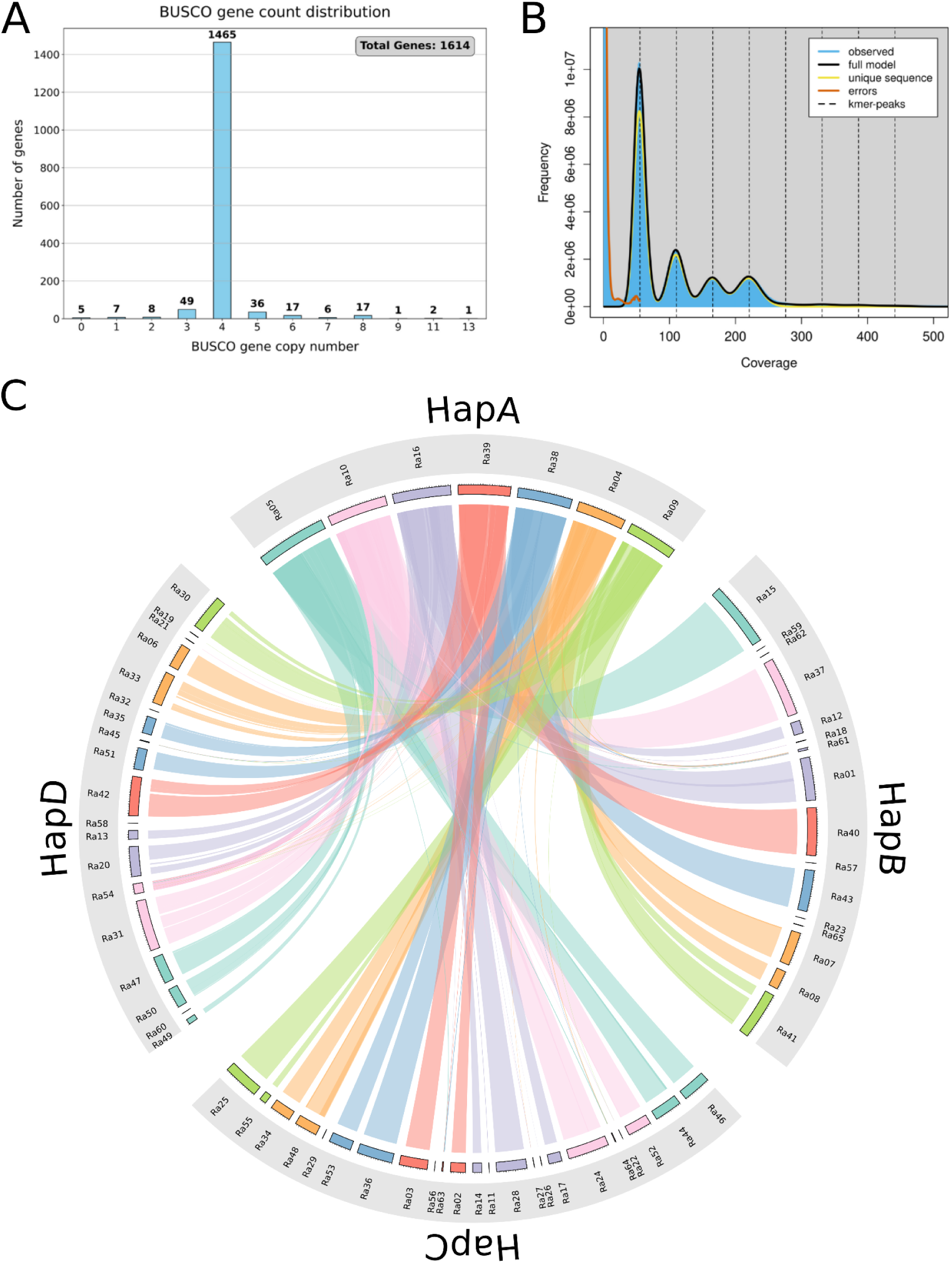
A: Visual representation of the amount of BUSCO genes against their abundance in the assembly. The graph is based on the BUSCO score of the RAS assembly utilizing the embryophyta_odb10 dataset. A full list of BUSCO genes and their respective copy number can be found in Table S2. B: Coverage K-mer spectra and fitted models for *R*. *armeniacus* using GenomeScope2. Note that the plot has four major peaks indicating tetraploidy C: shows an overall good homology between the different pseudohaplophases with links connecting the corresponding contigs with its reference on HapA. However, short contigs show links to several different contigs from HapA indicating, that these regions cannot be assigned with certainty to one or the other pseudohaplophases as they occur on several contigs within the central region of the reference chromosome, this could indicate that these regions are duplicated within the genome of *R. armeniacus* on different chromosomes. However, all short contigs were assigned to a pseudochromosome based on links.It should be noted that the accuracy of assignment to pseudohaplophases would benefit from additional external support.

Lastly, the RAS genome assembly was sorted into haplophases based on homology and scaffolding to its close relative *Rubus caesius.* This resulted in 4 pseudohaplophases with pseudohaplophase A (HapA) consisting of 7 pseudochromosomes (Table 2). The other haplophases were assigned based on decreasing contig length per mapping and increasing contig numbers from pseudohaplophase B to D (see Table S1).

**Table 2:** Statistics of the pseudohaplophases of *Rubus armeniacus* based the RAS assembly including the total number of contigs, N50 in Mbp, BUSCO score (C: complete, S: single copy, D: duplicated, F: fragmented, M: missing BUSCO genes) based on the rosaceae_odb12 dataset, and the LTR Assembly Index (LAI).

|  | HapA | HapB | HapC | HapD |
| --- | --- | --- | --- | --- |
| **assembly size [bp]** | 339174087 | 321326537 | 313912918 | 278927636 |
| **total contigs** | 7 | 16 | 23 | 19 |
| **N50 [mbp]** | 46 | 41 | 24 | 25 |
| **BUSCO score** | C:98.8%<br>[S:97.1%,D:1.8%]<br>,F:0.3%,M:0.9%,<br>n:10071 | C:94.7%<br>[S:90.2%,D:4.5%]<br>,F:0.3%,M:5.0%,<br>n:10071 | C:98.8%<br>[S:96.5%,D:2.3%],<br>F:0.3%,M:0.9%,<br>n:10071 | C:95.3%<br>[S:89.3%,D:6.0%],<br>F:0.2%,M:4.5%,<br>n:10071 |
| **LAI** | 24.72 | 26.97 | 24.69 | 27.15 |

All pseudohaplophases show a high completeness score with HapA and HapC displaying the highest completeness (98.8% of all BUSCO genes being represented), with only a low duplication rate of 1.8% and 2.3%, respectively. HapB and HapD display a slightly decreased completeness score with 94.7% and 95.3%, respectively, and increased duplication of 5% and 4.5%, respectively (Table 2).

The correct assignment of contigs to the different haplophases was supported and visualized utilizing a linkage map in the form of a Circos plot (Figure 2C).

### Functional and structural annotation

The structural annotation of *R. armeniacus* RAS revealed a total of 118,183 protein coding genes within the genome assembly. These can be assigned to different pseudohaplophases leading to 29054, 29079, 31349, and 28701 structurally annotated genes for the pseudohaplophases A, B, C, and D, respectively. Utilizing several functional annotation pipelines a total of 108,229 genes were functionally annotated (Pucker and Wolff, 2026).

The focus of this study is the induction and development of the colour change in the berries from green to black during ripening. This colour change has been attributed to a pigment class called anthocyanins (Patel and Rao, 2014; Sapers et al., 1986). In order to further analyse this topic, structural genes of the anthocyanin biosynthesis as well as the known transcription factors were annotated and further analysed in detail.

### Annotation of anthocyanin biosynthesis genes

The anthocyanin biosynthesis pathway is widely conserved across the angiosperms and has been intensely studied in a variety of species (Davies et al., 2024; Pucker and Selmar, 2022; Winkel-Shirley, 2001).

All relevant functional anthocyanin biosynthesis genes were identified and annotated, and are represented at least once within each pseudohaplophase (Table 3) further supporting the fully phased assembly.

**Table 3:**
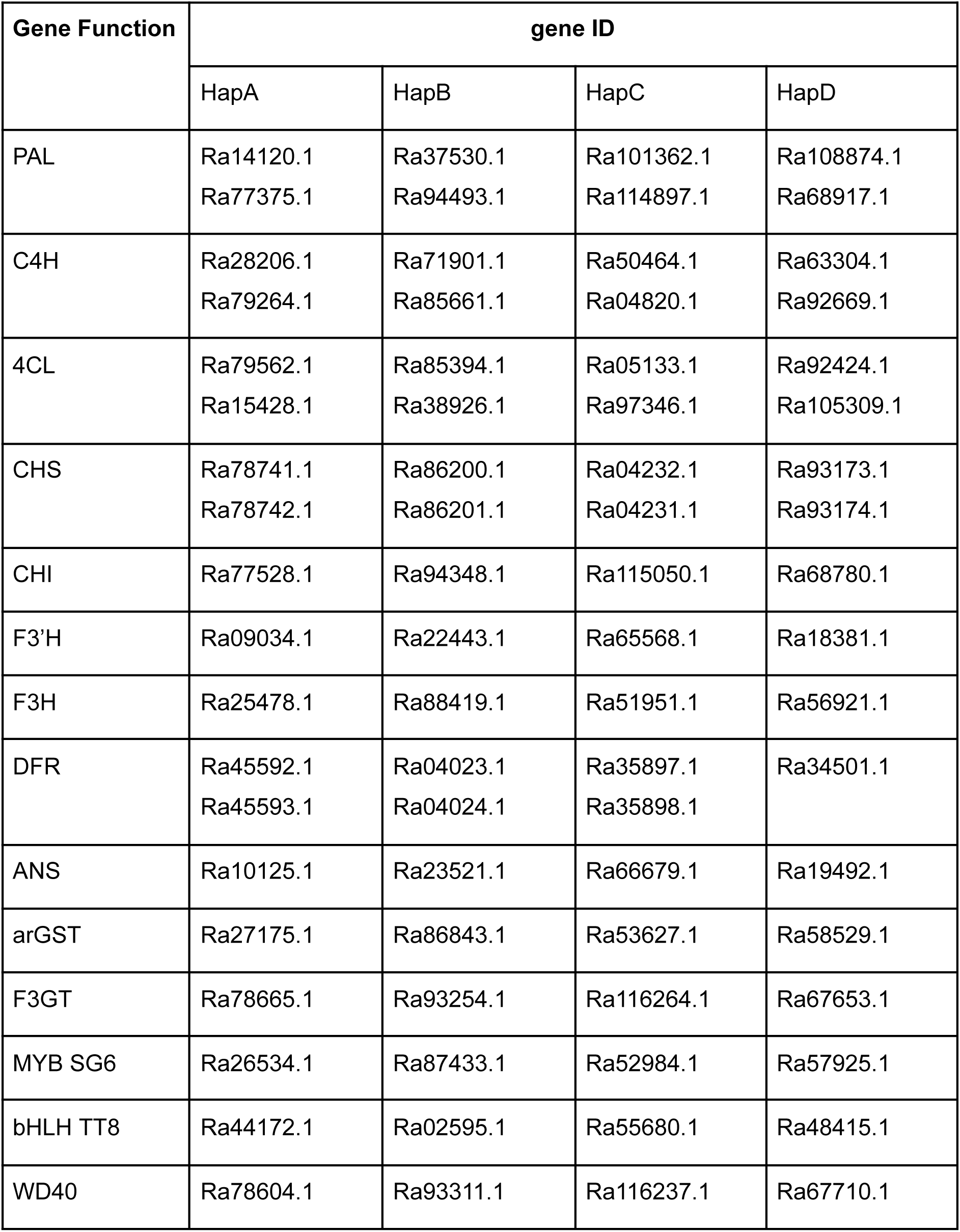
Overview of the distribution of anthocyanin biosynthesis genes and transcription factors across the different pseudohaplophases. Enzyme functions are annotated as follows: PAL (Phenylalanine ammonia lyase), C4H (Cinnamate 4-hydroxylase), 4CL (4-coumarate-CoA ligase), CHS (Naringenin-chalcone synthase), CHI (Chalcone isomerase), F3H (Flavanone 3-hydroxylase), F3’H (Flavonoid 3’-hydroxylase), F3’5’H (Flavonoid 3’,5’-hydroxylase), DFR (Dihydroflavonol 4-reductase), ANS (Anthocyanidin synthase), arGST (anthocyanin-related Glutathione S-transferase), F3GT (UDP-dependent anthocyanidin-3-O-glucosyltransferase).

| Gene Function | gene ID |  |  |  |
| --- | --- | --- | --- | --- |
|  | HapA | HapB | HapC | HapD |
| PAL | Ra14120.1<br>Ra77375.1 | Ra37530.1<br>Ra94493.1 | Ra101362.1<br>Ra114897.1 | Ra108874.1<br>Ra68917.1 |
| C4H | Ra28206.1<br>Ra79264.1 | Ra71901.1<br>Ra85661.1 | Ra50464.1<br>Ra04820.1 | Ra63304.1<br>Ra92669.1 |
| 4CL | Ra79562.1<br>Ra15428.1 | Ra85394.1<br>Ra38926.1 | Ra05133.1<br>Ra97346.1 | Ra92424.1<br>Ra105309.1 |
| CHS | Ra78741.1<br>Ra78742.1 | Ra86200.1<br>Ra86201.1 | Ra04232.1<br>Ra04231.1 | Ra93173.1<br>Ra93174.1 |
| CHI | Ra77528.1 | Ra94348.1 | Ra115050.1 | Ra68780.1 |
| F3'H | Ra09034.1 | Ra22443.1 | Ra65568.1 | Ra18381.1 |
| F3H | Ra25478.1 | Ra88419.1 | Ra51951.1 | Ra56921.1 |
| DFR | Ra45592.1<br>Ra45593.1 | Ra04023.1<br>Ra04024.1 | Ra35897.1<br>Ra35898.1 | Ra34501.1 |
| ANS | Ra10125.1 | Ra23521.1 | Ra66679.1 | Ra19492.1 |
| arGST | Ra27175.1 | Ra86843.1 | Ra53627.1 | Ra58529.1 |
| F3GT | Ra78665.1 | Ra93254.1 | Ra116264.1 | Ra67653.1 |
| MYB SG6 | Ra26534.1 | Ra87433.1 | Ra52984.1 | Ra57925.1 |
| bHLH TT8 | Ra44172.1 | Ra02595.1 | Ra55680.1 | Ra48415.1 |
| WD40 | Ra78604.1 | Ra93311.1 | Ra116237.1 | Ra67710.1 |

The majority of the anthocyanin biosynthesis genes are represented in the genome in copies of 4, one on each pseudohaplophase on the homologous pseudochromosomes. However, there appears to be two copies of the biosynthesis genes *PAL, C4H, 4CL,* and *CHS* which act upstream of the anthocyanin biosynthesis. Additionally, *DFR* is present in two copies in HapA, HapB, and HapC. While the duplications of *PAL*, *C4H,* and *4CL* are not on the same pseudochromosomes or contigs. The duplications of *CHS* and *DFR* appear to be tandem duplications in close proximity to each other in the genome. The duplication of *CHS* can be found in every haplophase while the tandem duplication of *DFR* is missing in HapD. This indicates that the duplication occurred prior to polyploidisation and was followed by a secondary loss of a *DFRb* copy within one haplophase of *R. armeniacus* The distribution of the anthocyanin biosynthesis genes further indicates the correct separation of the haplophases as well as the tetraploidy of *R. armeniacus*.

### Analysis of expression patterns and metabolic quantification of cyanidin-3-O-glucoside

Overall, the expression of the anthocyanin biosynthesis genes shows a good correlation across the haplophases, which is why the gene expression was summarised across the haplophases and the TPM values are represented as sums (Figure 3) .

**Figure 3:**
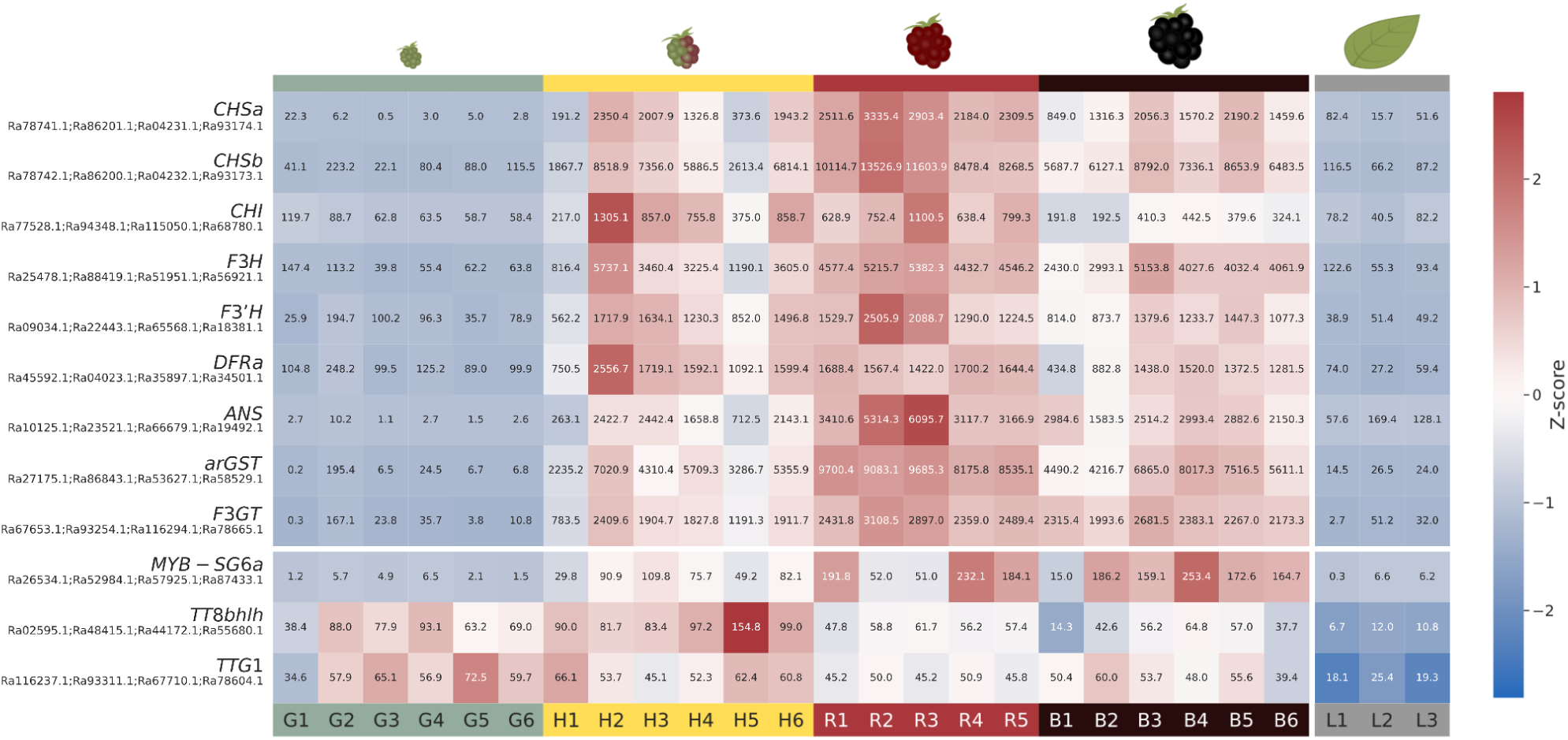
Heatmap showing expression of the genes of the anthocyanin biosynthesis pathway during different stages of fruit ripening: green (G), half-red (H), red (R), and black (B) with the numbers representing biological replicates. The expression is plotted as a sum of transcripts per million (numbers in tiles) per enzyme function and Z-score normalized over all samples per gene indicated by a diverging colour map. Individual tpm values can be found in Data S1. The labels include the functional label as well as the gene ID of the respective enzyme in the *R. armeniacus* annotation or transcription factor in the order hapA, hapB, hapC, and hapD. Additionally, the transcriptional machinery consists of subgroup 6 MYB (MYB-SG6), a bHLH factor (TT8 bHLH) and the WD40 Protein (TTG1). If more than one gene copy was found on any haplophase this is indicated with a or b.

The anthocyanin biosynthesis genes do not show high expression in the green berries (Figure 3). During the ripening from green to red, the anthocyanin biosynthesis is being activated and the expression of the anthocyanin biosynthesis genes is increasing.

*CHS* is present as a tandem duplication within the genome in all four haplophases, both copies of the gene show a very similar expression pattern, with low expression in green berries and an increase of expression over the course of development to red berries followed by a decrease in expression in the dark berries. *DFR* is also present as a tandem duplication, but the copies do not show the same expression pattern. Only one of the *DFR* copies (*DFRa*) shows significant expression, while the other copy (*DFRb*) does not show expression in any of the haplophases. The result of KIPEs indicates that DFRb possesses all relevant amino acid residues to be a functional enzyme (Table S3), indicating that the lack of gene expression might be due to a varying biological function or expression in a different part of the plant rather than a lack of biochemical functionality. Additionally, it has been shown that *DFR* displays a substrate-preference based on a specific residue at the position 133 relative to the *DFR* gene from *Arabidopsis thaliana (Choudhary and Pucker, 2024)*. For *R. armeniacus*, the expressed *DFRa* has the amino acid Asparagin (N) at the relevant position, which has been shown to produce mostly cyanidin (Johnson et al., 2001; Miosic et al., 2014), interestingly the unexpressed *DFRb,* contains the amino acid threonin (T) at the relevant position (see Figure S1), while this has not been associated with any substrate specificity, it has been identified as a functional DFR variant in several species including *Tacca chanteri* (Oliveira and Pucker, 2026), *Dioscorea alata,* and *Dioscorea rotundata (Choudhary and Pucker, 2024)*. This indicates that the identified copies of *DFRb*, might not represent a pseudogene but rather serve a specific function within *R. armeniacus* outside of the berry or leaf.

To further understand the accumulation of anthocyanins within the berries it is important to not only look at the functional pathway genes but to understand and analyse the regulatory factors controlling the expression. Anthocyanin regulation has been largely attributed to certain bHLH and MYB transcription factors in combination with the WD40 proteins. The expression of anthocyanin-regulating MYB and bHLH transcription factors is visualized in Figure 3 as well. TTG1 is steadily expressed throughout all ripening stages of the berries with low fluctuation between the lowest expression of 34.6 TPMs and the highest of 72.5 TPMs both occurring in the green berries. This distribution is to be expected, as TTG1 is not only involved in the regulation of anthocyanins but it serves a more diverse role within the plant including a range of functions such as trichome development, abiotic stress response, seed coat development, and root growth (Lim et al., 2022; Tan et al., 2021; Walker et al., 1999). MYB_SG6 is not active in the green berries and shows expression values of < 3 TPMs in green berries. With the ripening of the berries through the half-red stage over the red and lastly the black stage of the berry the expression of MYB_SG6 increases with the highest expression values reaching 253.4 TPMs in the black berries. In contrast, the *TT8* bHLH shows expression within the green berries and reaches the highest TPM values in the half-red stage of berry ripening. For both the red and black berries, a decreased expression of *TT8* bHLH can be observed.

The transcriptional analysis of *R. armeniacus* revealed the high expression of all relevant anthocyanin genes to produce cyanidin-based derivatives.

To further understand whether the dark pigmentation of *R. armeniacus* can be linked to specific anthocyanins or an overall increased accumulation of anthocyanins through the fruit ripening, a metabolic quantification of the anthocyanins was performed.

Firstly, a Thin Layer Chromatography (TLC) was utilized to verify that cyanidin-3-O-glucoside is the relevant anthocyanin present in blackberry fruit. Afterwards, the quantification of cyanidin-3-O-glucoside was performed using a HPLC-MS (Figure 4A, Figure S2).

**Figure 4:**
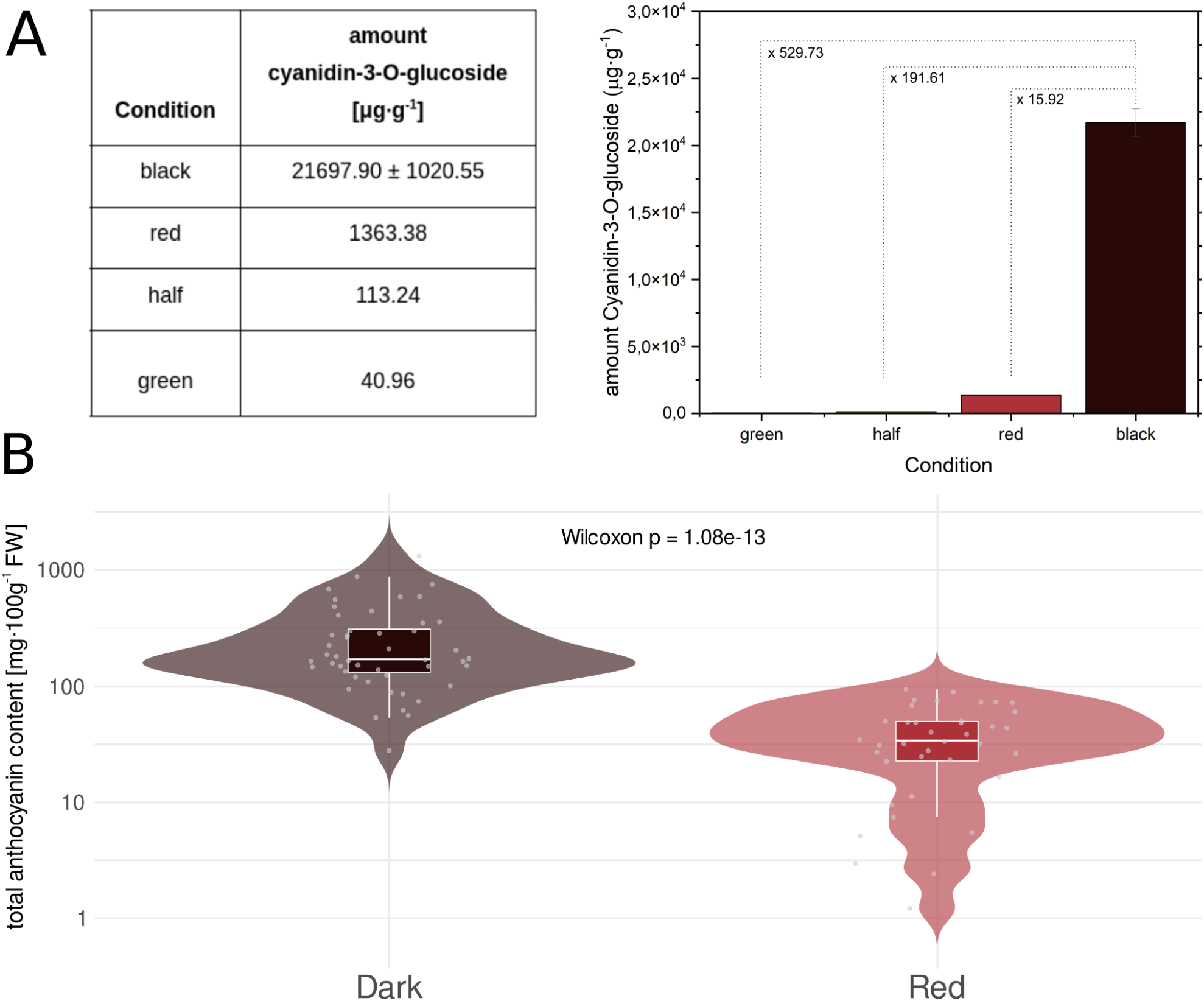
A: Content of cyanidin-3-O-glucoside in μg·g^-1^ in methanolic extract of pooled (n=3) berries of *R. armeniacus* at different ripening stages. For the black berry two different dilutions of methanolic extract were measured, the graph shows the mean of these two measurements with a calculated standard deviation as error bars. Additionally, the factor of anthocyanin content increase between black and the other conditions (dotted lines) is included in the graph. B: Distribution of total anthocyanin content in mg/100g fresh weight (FW) in different plant species with dark (n=48) and red (n=38) fruits based on publicly available datasets. A full list of included species can be found in Data S2-S4. The data is shown as violin plots with embedded boxplots (median and interquartile range) and individual observations (jittered points). Values are displayed on a log_10_-scaled y-axis. Statistical significance between groups was assessed using a Wilcoxon rank-sum test (p =1.08^-13^).

**Figure 5:**
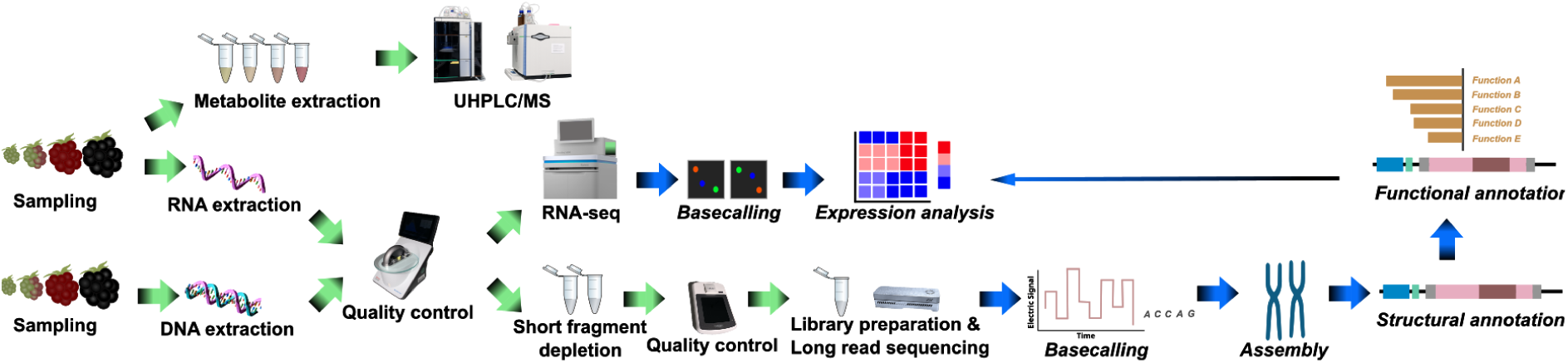
Multiomics workflow for the analysis of anthocyanin accumulation in *R. armeniacus*.

The extract of the unripe green berry shows a very low content of cyanidin-3-O-glucoside with 40.96 μg·g^-1^ dry weight. Throughout the ripening and with increasing intensity of colour the content of cyanidin-3-O-glucoside increases with 113.24 μg·g^-1^ in half red-half green berries, 1363.38 μg·g^-1^ in red berries and reaching the highest mean concentration of cyanidin-3-O-glucoside of 21697,90 μg·g^-1^, making up slightly above 2% of berry dry weight. This represents a 16-fold increase in cyanidin-3-O-glucoside in comparison to the red berries.

## Discussion

### Comparison of Genomic Assembly quality

To the best of our knowledge, there are only five genome sequences available which represent *Rubus* species with darkly pigmented fruits, while the other 11 available genome sequences are of species producing red fruits, highlighting the need for further genetic resources regarding the dark pigmentation of *Rubus*. Most available genome sequences, with the exception of *Rubus hochstetterorum,* show a high level of completeness with BUSCO scores ranging from 97.5% for *Rubus parviflorus* to 99.1% for *Rubus caesius* (see Table S4). Similarly, the genome assembly of *Rubus armeniacus* presented in this study shows comparable BUSCO scores reaching a completeness BUSCO score of 99.1% for the whole genome sequence and 98.8% for haplophase A. All available genomes displayed a high completeness value based on BUSCO score, however the BUSCO score alone does not allow for a meaningful comparison or distinction based on the assemblies quality. More importantly, while the completeness of all assemblies is high, the number of contigs representing the genome varies greatly between the assemblies available for different species. *Rubus* species harbour chromosomes and range from diploid to tetrapoloid species. The lowest number of contigs or scaffolds for a haploid assembly is 11 scaffolds for the diploid species *Rubus idaeus Malling Jewel* (Price et al., 2023). Due to the high completeness with low duplication this assembly is assumed to be a haploid genome sequence. *Rubus alceifolius* is represented by a complete genome-level assembly including 14 scaffolds and a BUSCO score of 99.0% with a duplication score of 88.3%, indicating 2x7 chromosome representation within this assembly (Zheng, 2025). The genome assembly of *Rubus armeniacus* consists of 65 contigs, without any scaffolding, and represents the tetraploid genome, when separated into pseudohaplophases, HapA consists of 7 chromosome-scale contigs, without a significant loss in completeness, indicating a high level of continuity for the assembly. Further the LAI score was generated for all available genome sequences (see Table S4) to allow for comparison of genome assembly quality across different assemblers and sequencing methods (Ou et al., 2018). The LAI scores for the available genome sequences of the *Rubus* genus stretch from 4.83 to 13.58. In general, the LAI score assigns assemblies into categories of draft (LAI < 10), reference (10 <= LAI < 20), and gold (20 <= LAI) assembly quality (Mokhtar et al., 2023; Ou et al., 2018). For the analysed 12 available assemblies, 11 fall into the category of draft assembly with LAI scores below 10, only the *Rubus alceifolius* falls into the category of reference assembly with a LAI score of 13.58. In contrast the whole genome sequence of *Rubus armeniacus* shows a much higher LAI score for both the whole assembly with 19.54 placing it at the higher end of reference assembly and 24.72 for hapA, indicating a gold quality assembly. Similarly high LAI scores, of 24.89, have been achieved for the diploid species of *Rosa sterilis*, a representative of the Rosaceae family (Zong et al., 2024). A large study analyzing 1136 diploid plant genome sequences assessed the quality based on LAI score revealed that out of those only 135 genome sequences were classified as gold quality, while 476 genome sequences classified as draft and 472 as reference (Mokhtar et al., 2023).

The assembly of *Rubus armeniacus* presented in this study consists of a total of 65 contigs without any additional scaffolding and is able to represent a single haplophase of the genome in chromosome scale contigs represented by 7 contigs, indicating a high level of continuity which is also indicated by a high N50 of 46 Mbp for only HapA and a N50 of 35.5 Mbp for the whole assembly. Due to the assembly’s high completeness and continuity, it not only provides a valuable addition to the genomic resources of the *Rubus* genus, it also provides the ideal basis for a comprehensive functional and structural annotation to further understand the underlying genetic functions at play within this genus, as well being a valuable resource to further study the genome evolution of the genus.

### The underlying mechanism of dark pigmentation in *R. armeniacus*

The expression pattern of the anthocyanin biosynthesis genes shows an increase in expression of the relevant genes during fruit ripening potentially explaining the colour shift from green to red, which has been supported by the accumulation of high concentrations of cyanidin-3-O-glucoside within the dark berry extract of *R. armeniacus*. While cyanidin-3-O-glucoside has been reported to be the most abundant anthocyanin within the dark fruited berries of *R. armeniacus*, the main anthocyanin present in *Rubus occidentalis*, the black raspberry has been shown to be cyanidin-3-O-rutinoside (Tian et al., 2006; Tulio et al., 2008), indicating that the mechanism for dark colour within the *Rubus* genus might not be conserved.

Similar observations have been reported for berries from other families such as *Morus atropurpurea Roxb.,* commonly known as mulberry (Huang et al., 2020). In a comparison of the gene expression level of key anthocyanin biosynthesis genes for the highly pigmented variety of *Morus atropurpurea Roxb* Da 10 with the white fruited variety Baisang (*Morus alba L*), it was revealed that the white variety showed significantly lower expression of all specialized anthocyanin biosynthesis genes, however an increase in the expression of the flavonol synthase was observed, hinting at an increased flux towards the colourless flavonols potentially explaining the observed phenotype (Huang et al., 2020). Similarly, a study of *Solanum nigrum*, identified the R2R3-MYB transcription factor *SnAN2* as a key regulator involved in the formation of dark berries, and demonstrated loss of dark pigmentation in berries for SnAN2 knockout mutants, which were only able to produce green berries (Heo et al., 2022).

Accumulation of anthocyanins is often associated with high expression of associated genes or the overexpression of certain transcription factors. For example, the increase in expression of *AmMYB10* has been shown to be the cause of anthocyanin biosynthesis induction and accumulation within *Aronia melanocarpa* during fruit ripening, a member of the Rosaceae family (Mahoney et al., 2022). Further, the anthocyanin biosynthesis was reactivated within *Solanum lycopersicum*, by the introduction of the bHLH protein Delila (AmDel) and the MYB protein Rosea1 (AmRos1) into a wild-type tomato, resulting in a dark phenotype, based on the activation of several enzymes involved in the anthocyanin biosynthesis (Butelli et al., 2008). This accumulation of anthocyanins could be further increased by the addition of a MYB12 transcription factor, highlighting the key role of transcriptional regulation within the development of dark phenotypes (Butelli et al., 2021).

In agreement with the previous studies, we observe an increased expression through the fruit ripening of *Rubus armeniacus* of the associated anthocyanin biosynthesis genes and a MYB-SG6 transcription factor, which is believed to activate the anthocyanin biosynthesis.

The metabolomic analysis revealed a substantial increase in anthocyanins during the ripening of the fruit, with cyanidin-3-O-glucoside accounting for over 2% of berry dry weight. Similarly high values of anthocyanins have been reported for other species of dark fruit and berries. A study on 20 different wild billberry (*Vaccinium myrtillus L.*) populations revealed cyanidin levels ranging between 6520 and 12140 μg·g^-1^ DW, depending on the location of the population of blueberries (Lätti et al., 2008). However, while in *R. armeniacus* only cyanidin-3-O-glucoside was identified as the main anthocyanin, *V. myrtillus* contains a variety of different anthocyanin derivatives. Delphinidine based anthocyanins reach similarly high levels as cyanidin based derivatives, while petunidin, malvidin, and pelargonidin were identified in lower quantities. The total anthocyanin content was identified in a range between 19340 and 38770 μg·g^-1^ DW for *V. myrtillus*, showing a higher content of anthocyanins then the values reported here for *R. armeniacus* (Lätti et al., 2008).

Further, the development of the anthocyanin biosynthesis was studied in *Aronia melanocarpa*, revealing the accumulation of cyanidin derivatives during the fruit ripening with cyanidin-3-O-galactoside appearing as the main anthocyanins (Mahoney et al., 2022).

Depending on the location of the *A. melanocarpa* cultivar the content of cyanidin derivatives ranged between 23303 and 55284 μg·g^-1^ DW in the final stage of ripening (Mahoney et al., 2022), showing a higher cyanidin content than *R. armeniacus*. The first stage of ripening of *A. melanocarpa* is defined as a dark red berry, similar in colour to the “R” stage presented for *R. armeniacus* in this study, shows much lower concentrations of cyanidin based anthocyanins present ranging from 4062 to 9379 μg·g^-1^ DW (Mahoney et al., 2022), placing it within range of the anthocyanin content of the red berry of *R. armeniacus*.

Further anthocyanin quantifications of red fruited plants (n=38) and plants with dark fruits (n=48) from publicly available databases (Haytowitz et al., 2018; Rothwell et al., 2016) were analysed. Revealing a significant difference in anthocyanin content between the red and dark fruited plants, with a median total anthocyanin content of 33.99 mg/100g FW for red fruits and 170.88 mg/100g FW for dark fruited plants (Data S4). This indicates that a certain threshold of anthocyanin accumulation must be overcome to facilitate the change of appearance from red to dark. However, in-depth analyses need to be performed to pinpoint a specific cutoff value, as well as analyse the underlying cause of this high accumulation in darkly pigmented fruit. Additionally, species specific factors would further influence the threshold and should be taken into consideration.

## Material and Methods

### DNA extraction and genome sequencing

High molecular genomic DNA was extracted based on a modified CTAB-method (Pucker, 2020; Siadjeu et al., 2020) from young leaves of *Rubus armeniacus* located in Braunschweig, Germany (DE-0-BONN-49806). The fresh plant material, consisting of young leaves, was homogenized with liquid nitrogen and taken up in CTAB1 buffer for resuspension. Afterwards, the resuspension was incubated for 30 minutes at 75°C while tubes were frequently inverted. After reaching room temperature, dichloromethane was added and mixed carefully. A 30 minute centrifugation step at room temperature followed with subsequent transfer of colorless upper phase and addition of CTAB2 buffer. The pellet was re-dissolved in 1M NaCl and participated with 100% isopropanol. After a washing step with 70% ethanol, the pellet was dissolved in CTAB-TE and incubated with RNase at room temperature overnight. For gDNA assessment, NanoDrop (Thermo Fisher Scientific) measurement, agarose gel electrophoresis, and Qubit (Thermo Fisher Scientific) measurements were conducted. To enrich long DNA fragments, a short read eliminator kit (SRE, Pacific Biosciences) was used with subsequent quantification with Qubit.

Library preparation was executed with 1µg gDNA according to the SQK-LSK109 protocol (Oxford Nanopore Technologies, ONT). Sequencing was performed on a MinION Mk1B (ONT) using a R9.4.1 flow cell.

Additional sequencing runs were performed with R10.4.1 flow cells. The library preparation was performed according to the SQK-LSK114 protocol with a DNA input of 1 ug.

Basecalling was conducted with dorado v0.8.3 (ONT) using the model dna_r10.4.1_e8.2_400bps_sup@v5.0.0 and detection of modified bases (--modified-bases “5mCG_5hmCG”) was performed, followed by an additional correction of the reads with HERRO v1 (Stanojević et al., 2024).

### RNA extraction and RNA-seq

For RNA extraction berries in four different growth stages from a *Rubus armeniacus* plant located in Braunschweig, Germany (52.28031563395348, 10.549554215883669), were collected and immediately frozen in liquid nitrogen and stored at -70 °C until sample preparation.

The berries were selected based on colour with the stages including: green (G), half-red (H), red (R), and black (B).

RNA extraction was performed according to (K. Wang et al., 2021), with the notable change that the extraction buffer was prepared according to (Reid et al., 2006), without the addition of spermidin and can be accessed at protocols.io (“Phenol-free RNA extraction of *Rubus armeniacus* berries”). The berries were removed from -70 °C storage and kept frozen on liquid nitrogen. For homogenization, one berry was ground in liquid nitrogen with mortar and pestle and transferred into two 2 mL reaction tubes containing the preheated extraction buffer. Followed by an incubation at 65 °C for 20 min during which the samples were inverted regularly. After the incubation the samples were centrifuged for 10 min with 11,000 g at 4 °C. The supernatant was transferred to a new 2 mL reaction tube and extracted with equal volume of chloroform:Isoamylalcohol (24:1) and centrifuged at 12,000 g for 10 minutes and the supernatant transferred to a new 2 mL reaction tube. The extraction was repeated. To precipitate the RNA, 0.25 volume of 10 M LiCl was added and gently mixed and incubated at 4 °C overnight. After a centrifugation at 12,000 g for 20 min at 4°C, the supernatant was discarded and the pellet was washed with 500 µL of ice cold ethanol. A centrifugation at 12,000 g at 4 °C for 5 minutes was carried out and the ethanol decanted. The resulting RNA pellet was dissolved in 100 µL TE buffer and precipitated with 300 µL cold 100% ethanol. After 5 min of centrifugation at 12,000 g at 4°C the supernatant was discarded and the pellet dissolved in 20 µL TE buffer.

The concentration of RNA was determined based on NanoDrop measurement and the integrity was assessed with agarose gel electrophoresis.

The RNA was stored at -70 °C until it was sent for RNA-seq. Paired-end RNA-seq (2 x 150 nt) was conducted using Illumina NovaSeq 6000.

### Genome sequence assembly

Three different assembly strategies were applied utilizing Shasta v0.14.0 (Shafin et al., 2020), Hifiasm v0.25.0 (Cheng et al., 2024) and NextDenovo v2.5.2 (Hu et al., 2024). For NextDenovo the estimated genome size was set at the expected tetraploid genome size of 1.5 Gbp and a minimum read length of 10 kbp. While Shasta was run using the Nanopore-r10.4.1_e8.2-400bps_sup-Herro-Jan2025 configuration file. Hifiasm was run with the --n-hap 4 flag to account for the tetraploid genome of *R. armeniacus*.

All contigs below 50 kbp were discarded and the statistics were determined utilizing the python script contig_stats3.py (de Oliveira et al., 2026).

The correctness of the assembly was assessed based on BUSCO v5.8.2 (Manni et al., 2021), with the dataset rosaceae_odb12, as well as the LTR Assembly Index, with default parameters and the size parameter set to 50000 (Ou et al., 2018).

### Separation of pseudohaplophases

To assess the ploidy of the generated *R. armeniacus* genome assembly, a custom Python script was used to analyse the copy number distribution of BUSCO genes (https://github.com/PuckerLab/Rubus_armeniacus/blob/main/busco_cov_hist_2.py). A plot was generated showing the number of BUSCO genes relative to their copy number present in the genome sequence.

To separate the haplophases, the genome sequences were first mapped against the reference pseudochromosomes of *Rubus caesius* using the MUMmer 3 module NUCmer (NUCleotide MUMmer) (Kurtz et al., 2004) with default settings. The resulting alignments were processed using the show-coords module with the parameters -TH rubus_v5_rc_hap1.delta, followed by sorting based on the alignment length (sort -k5,5nr). The output was further processed with three custom python scripts. The first script (nucmer_process_rc.py) was used to merge mappings of the same contig that were in the proximity of 1000 bp into a single mapping. The second script (find_nucmer_dups_summary.py) reformatted the resulting merged mapping and identified any regions where different contigs mapped to the same region of the reference assembly. Lastly, the third python script (plot_single_four.py) generated a graphical representation of the merged mappings for each reference chromosome (https://github.com/PuckerLab/Rubus_armeniacus). For each chromosome, two plots were created: one which shows all mappings with any number of duplications of the different contigs on the reference pseudochromosome and a second graph which only shows mapping regions where four different contigs overlap (Figure S3).

Overlapping contigs were manually curated into four different pseudohaplophases, with the largest contigs being sorted into HapA. The remaining contigs were assigned to the pseudohaplophases based on descending contig length.

Additionally, RagTag v2.1.0 with default parameters (Alonge et al., 2022) was utilized to place shorter contigs and fill any gaps still present in the pseudohaplophases. The genome sequence of *R. caesius* (GCA_964235055.1, Darwin Tree of Life) was used as a reference assembly. To ensure haplophase-specific scaffolding, contigs assigned to other pseudohaplophases were excluded using the -exclude option of RagTag. Short contigs were iteratively assigned to HapB, HapC, and HapD based on best match, with each pseudohaplophase scaffolded independently. The resulting mappings were used to order contigs within each pseudohaplophase.

The correctness of the pseudohaplophases was assessed based on BUSCO v5.8.2 (Manni et al., 2021), with the dataset rosaceae_odb12, as well as the LTR Assembly Index, with default parameters and the size parameter set to 50000 (Ou et al., 2018).

### Structural and functional annotation

A homology-based gene prediction of the *R. armeniacus* genome sequence was performed using GeMoMa v1.9 (Keilwagen et al., 2018, 2016), based on annotations of seven closely related *Rubus* species namely *Rubus idaeus* Autumn Bliss (Price et al., 2023), *Rubus argutus* Hillquist (Brůna et al., 2023), *Rubus chingii* (L. Wang et al., 2021)*, Rubus idaeus* Anitra (Davik et al., 2022), *Rubus idaeus* Joan J (Zhou et al., 2023), *Rubus Idaeus* Malling Jewel (Price et al., 2023) and *Rubus occidentalis* (VanBuren et al., 2018). The resulting seven gene annotation sets were filtered and merged using GeMoMa Annotation Filter (GAF) with parameters f="start==’M’ and stop==’*’ and (isNaN(score) or score/aa>=4.5) and aa>50" atf="tie==1 or sumWeight>1". BUSCO v5.8.2 with the rosaceae_odb12 dataset was used to assess the completeness of the annotations.

Additionally, a structural gene annotation was performed using BRAKER3 (v3.0.8) (Gabriel et al., 2024) to predict protein-coding genes, incorporating RNA-seq data as external evidence. Total RNA was extracted from blackberry (*Rubus* spp.) fruit tissues at multiple developmental stages and reverse transcripted into cDNA for sequencing with Illumina paired-end technology (see “RNA extraction and RNA-seq” for details). High-quality RNA-seq reads were aligned to the reference genome sequence using STAR v2.7.11b with standard parameters (Dobin et al., 2013). The resulting BAM file was used as input for BRAKER v3.0.8 (Gabriel et al., 2024) as RNA-seq hint evidence. BRAKER3 was executed with default parameters, utilizing the STAR-aligned RNA-seq data to generate extrinsic evidence.

BRAKER3 produces an annotation in GFF3 format. However, in order to merge and filter the GFF3 file in the same manner as the GeMoMa annotation, attributes which define the quality of the gene models, such as RNA-seq coverage and intron information from RNA-seq alignments had to be added to the BRAKER3 annotation. A coverage.bedgraph and introns.gff were generated utilizing GeMoMa’s ERE module based on the RNA-seq mapping file in BAM format. These files are used to add quality attributes to gene models based on expression data, using GeMoMa CLI AnnotationEvidence, with the parameters c=UNSTRANDED.

The resulting annotation_with_attributes.gff file, includes attributes such as average RNA-seq coverage per gene (avgCov), fraction of transcript’s introns supported by RNA-seq (tie), amino acid sequence length (aa). The resulting annotation GFF file was filtered and merged with the GeMoMa GFF file with GeMoMa Annotation Filter (GAF) with parameters f="start==’M’ and stop==’*’ and (isNaN(score) or score/aa>=4.5) and aa>50" atf="tie==1 or sumWeight>1". BUSCO v5.8.2 with the rosaceae_odb12 dataset was used to assess the completeness of the annotations.

Several approaches were taken to generate a functional annotation. First a functional annotation was performed using InterProScan v5.75-106.0 (Jones et al., 2014), as well as using the Funannotate v1.8.17 “annotate” function (Palmer and Stajich, 2020). Additionally, a functional annotation was generated based on similarity to *Arabidopsis* genes utilizing the Python script construct_anno.py (Pucker and Iorizzo, 2023). Lastly, the three annotations were merged into one csv file.

To identify the genes involved in the anthocyanin biosynthesis KIPEs v0.35 (Rempel et al., 2023) was used, utilizing the FlavonoidBioSynBaits_v3.3 and the results were filtered to only include gene candidates including 100% of conserved amino acid residues. Additionally, regulatory genes such as bHLH and MYB transcription factors were identified utilizing bHLH_annotator v1.04 (Thoben and Pucker, 2023) and MYB_annotator v1.0.1 (Pucker, 2022), respectively. To further confirm the identified MYBs and bHLH and identify WD40 candidates the collect_best_blast.py script (Pucker and Iorizzo, 2023) was utilized to identify candidates based on their similarity to *Malus domestica*. The identified candidates were mapped to respective bait peptide sequences of bHLH and MYB reference sequences from a variety of species according to (Choudhary et al., 2025) using MAFFT v7.505 (G-INS-i algorithm) (Katoh et al., 2019; Kuraku et al., 2013). The resulting mapping was translated to codon sequences using pxaa2cdn, part of phyx v1.3.2 (Brown et al., 2017). Lastly the respective phylogenetic tree was calculated for MYBs, bHLH and WD40 identification, using IQ-TREE v2.0.7 with the parameters -B 1000 -m MFP, the best-fit model according to BIC: GTR+F+R9, TIM3+F+R4 and JTT+R3 were identified by ModelFinder for the MYB, bHLH and WD40 tree generation, respectively.

### Expression analysis of anthocyanin biosynthesis genes and transcription factors

To quantify transcript abundance based on the coding sequences kallisto v0.51.1 (Bray et al., 2016) was utilized. A previously developed Python script kallisto_pipeline3.py (Pucker and Iorizzo, 2023) was used to run kallisto across all input files based on berry colour and the resulting count tables were merged with the Python script merge_kallisto_output3.py (Pucker and Iorizzo, 2023).

The countables were filtered for the geneIDs associated with the anthocyanin biosynthesis genes, bHLH and MYB transcription factors, respectively. A customised Python script was utilized to create heatmaps visualising the expression of the genes across the four haplophases by addition of the TPM values for each gene (https://github.com/PuckerLab/Rubus_armeniacus/blob/main/plot_single_four.py). To compare the genes across the different samples a z-score normalization was performed.

### Metabolomic analysis of cyanidin-3-O-glucoside from metabolic berry extracts

The fruits of *Rubus armeniacus* (Braunschweig, Germany (52.28031563395348, 10.549554215883669)) were freeze dried overnight and and for each ripening stage, green, half-red-half-green, red, and black, three berries were pooled and ground by mortal and pester into a paste. For the metabolic extraction of anthocyanins 200 mg of each berry pool were measured and 1 mL of HPLC-grade MeOH was added. Sonification in an ultrasound bath was performed for 15 minutes and afterwards the samples were centrifuged at 20,000 g for 10 minutes. The clear supernatant was transferred to a clean reaction tube and utilized for the following quantification of cyanidin-3-O-glucoside. The following dilutions were used for the different berry extracts: green 1:5, half-red-half-green 1:10, red 1:20 and black was measured with two dilutions 1:200 and 1:500.

High-resolution accurate-mass (HRAM) measurements of standard and extract solutions were performed using a Vanish Horizon ultra-high-performance liquid chromatography (UHPLC) system from Thermo Fisher Scientific, comprising a VH-P10-A pump, a VH-A10-A autosampler and a VH-C10-A column oven (not temperature-controlled), coupled to an Exploris 120 HRMS Orbitrap instrument (Thermo Fisher Scientific, Waltham, USA). The standards of Cyanidin-3-O-glucosid and naringenin were obtained from Carl Roth (Karlsruhe, Germany). The MS method included the following settings: H-ESI spray voltage (positive) of 3,500 V; ion transfer tube temperature and vaporizer temperature of 325 °C and 350 °C, respectively. The sheath gas, aux gas and sweep gas were set to 50, 10 and 1 arb. units, respectively. Full scan HRAM data were recorded for the m/z range of 250–950 at an Orbitrap resolution of 120,000. Prior to each UHPLC-MS run, a one-point mass calibration was performed using the built in EASY-IC.

Chromatographic separation was achieved using a Discovery HS F5 analytical column 150 mm × 2.1 mm, 3 µm (Supelco, Bellefonte, USA). Mobile phase A consisted of water containing 1% formic acid, and mobile phase B consisted of methanol containing 1% formic acid. The flow rate was 0.300 mL/min, with a chosen injection volume of 2 μL. The separation gradient was 5% B (0–2 min), followed by a gradient to 100% B from 2 to 22 min. The conditions were then maintained at 100% B (22-25 min), before changing back to 5% B (25–26 min). A final re-equilibration step was performed at 5% B (26–36 min).

Data acquisition, evaluation and quantification were performed using Thermo Scientific Xcalibur Version 4.6.67.17, Thermo FreeStyle 1.8 SP2, and Thermo Xcalibur Qual Browser 4.4.67.17. A four-point calibration curve ranging from 1 to 100 µg/mL was established using reconstructed ion chromatograms (RICs) for cyanidin-3-O-glucoside and naringenin at m/z 449.10784 and 273.07575, respectively, with a mass tolerance of ±2.5 ppm. This corresponds to the charged ions of cyanidin-3-O-glucoside ([C_21_H_21_O_11_]^+^) and the protonated pseudo-molecular ion of naringenin ([C_15_H_12_O_5_+H]^+^).

### Comparison of publicly available metabolomics data for red and dark fruits

Publicly available quantification data of anthocyanins for different fruits, as mg/100g FW, was collected from PhenolExplorer (Rothwell et al., 2013) as well as USDA Database for the Flavonoid Content of Selected Foods (Bhagwat and Haytowitz, 2018). The data was filtered to only contain raw or frozen fruits and all processed food items such as juices or powders were removed. For data extracted from PhenolExplorer, the inbuilt function to extract a total anthocyanin content for each species was utilized, while the total anthocyanin content was calculated for the database downloaded from USDA Database for the Flavonoid Content of Selected Foods.

The samples were grouped into the categories red and dark based on their fruit colours and an R script was utilized to perform a Wilcoxon rank sum test with continuity correction to determine the significance of the difference in median anthocyanin content based on colour. The exact data matrix used for the analysis can be found in Data S4. Lastly, the data was visualised as violin plots with embedded boxplots (median and interquartile range) and the individual sample values are shown as jittered points. Values are displayed on a log₁₀-scaled y-axis.

## Declarations

### Ethics approval and consent to participate

Not applicable

### Consent for publication

Not applicable

### Conflict of Interest Statement

Not applicable.

### Funding

Not applicable

### Authors’ contributions

BP and KW planned and designed the study. KW, MN and CT performed DNA extraction and sequencing. KW, MN and CT performed the bioinformatic analysis. KW performed the RNA and metabolite extraction. TB performed the HRAM and UHPLC and analysis.

KW interpreted the results, and wrote the manuscript.All authors have read the final version of the manuscript and approved its submission.

## Acknowledgements

This work was supported by the de.NBI Cloud within the German Network for Bioinformatics Infrastructure (de.NBI) and ELIXIR-DE (Forschungszentrum Jülich and W-de.NBI-001, W-de.NBI-004, W-de.NBI-008, W-de.NBI-010, W-de.NBI-013, W-de.NBI-014, W-de.NBI-016, W-de.NBI-022). The work was partially funded by the Deutsche Forschungsgemeinschaft (DFG, German Research Foundation) project number 511817356.

We thank all members of the Plant Biotechnology and Bioinformatics group for their support and feedback during the process. We are grateful for the excellent support provided by the team of the University of Bonn Botanic Gardens.

## Data availability statement

All custom scripts utilized for the analysis of the data are available at https://github.com/PuckerLab/Rubus_armeniacus.

The genome sequence and annotation associated with this publication can be accessed at https://doi.org/doi:10.60507/FK2/L7LG2L

All sequencing reads for this project, both RNA-seq and DNA reads, are available at ENA under PRJEB63972.

## Supporting Information

### Supporting figures

**Figure S1:** Mafft alignment of the DFR candidates identified in *Rubus armeniacus*, highlighting the difference in amino acid residue responsible for substrate preference

**Figure S2**: HPLC separation and HRAM MS of the different berry extracts, in comparison to the Cyanidin-3-o-glucoside reference

**Figure S3:** Mappings of the contigs of *Rubus armeniacus* against the reference chromosomes of *Rubus caesius*. The first plot shows mappings with any number of duplications of the different contigs on the reference pseudochromosome and a second graph which only shows mapping regions where four different contigs overlap

### Supporting tables

**Table S1:** Overview of assignment of contigs to the different pseudohaplophases and their respective contig size in bp.

**Table S2:** Full table of identified BUSCO genes in the genome and their localisation on the contigs. BUSCO genes identified in multiples are listed as Duplicated.

**Table S3:** All identified structural genes involved in the flavonoid biosynthesis which contain 100% of conserved amino acid residues relevant for the function of the enzyme in comparison to the KIPEs bait sequences.

**Table S4:** Comparison of available genome sequences for the genus *Rubus* with an assembly level of scaffold or higher.

### Supporting data

**Data S1:** TPM values of anthocyanin biosynthesis genes across the different berry ripening stages as well as leaf tissue.

**Data S2:** All data included in the analysis of anthocyanin content in red vs dark fruits/berries extracted from USDA Database for the Flavonoid Content of Selected Foods and their respective original source.

**Data S3:** All data included in the analysis of anthocyanin content in red vs dark fruits/berries extracted from PhenolExplorer and their respective original source.

**Data S4:** Dataframe including the species name, total anthocyanin content and assignment to red or dark used for the analysis of difference in anthocyanin content between red and dark berries.

## Notes

https://github.com/PuckerLab/Rubus_armeniacus

https://doi.org/10.60507/FK2/L7LG2L

### Competing Interest Statement

The authors have declared no competing interest.

