## Supplementary material for "*Rubus armeniacus* genome sequence reveals the secrets of blackberry anthocyanin biosynthesis": Data S4

| | | |
|---|---|---|
| 53,64 | Acai. berries. purple. fresh | Dark |
| 61,94 | Acai. berries. purple. frozen | Dark |
| 204,87 | Acai. fruit pulp/skin. powder | Dark |
| 22,55 | Acerola. (west indian cherry). raw | Red |
| 75,79 | Aestivalis grape [Black] | Red |
| 49,89 | American cranberry | Red |
| 32 | American cranberry | Red |
| 49,89 | American cranberry | Red |
| 5,5 | Apple. skin only | Red |
| 1,22 | Apples. Gala. raw. with skin | Red |
| 2,97 | Apples. Red Delicious. raw. without skin | Red |
| 89 | Arctic bramble berries | Red |
| 299 | Bilberry, raw | Dark |
| 285,21 | Bilberry. raw | Dark |
| 878,12 | Black chokeberry | Dark |
| 444,25 | Black chokeberry | Dark |
| 1316,65 | Black elderberry | Dark |
| 163,9 | Black huckleberry | Dark |
| 589 | Black raspberry, raw | Dark |
| 100,61 | Blackberries. raw | Dark |
| 146,8 | Blackberry, raw | Dark |
| 172,59 | Blackberry. raw | Dark |
| 225,04 | Blackcurrant, raw | Dark |
| 592,23 | Blackcurrant. raw | Dark |
| 163,3 | Blueberries. cultivated (highbush). raw | Dark |
| 94,25 | Blueberries. frozen. unsweetened | Dark |
| 148,61 | Blueberries. rabbiteye. raw | Dark |
| 150,26 | Blueberries. wild (lowbush). raw | Dark |
| 124,5 | Bog bilberry, raw | Dark |
| 273 | Buffalo | Dark |
| 209,95 | Cabbage. red. raw | Dark |
| 298 | Canada blueberry | Dark |
| 151,75 | Cascade huckleberry | Dark |
| 27,82 | Cedar bay cherry. raw | Red |
| 2,42 | Cherries. sour. dry. sweetened | Red |
| 7,45 | Cherries. sour. dry. unsweetened | Red |
| 34,53 | Cherries. sour. powder | Red |
| 11,24 | Cherries. sour. red. frozen. unsweetened | Red |
| 33,44 | Cherries. sour. red. raw | Red |
| 31,98 | Cherries. sweet. raw | Red |
| 349,79 | Chokeberry. raw | Dark |
| 262,49 | Cowpeas. black seed cultivar. mature seeds. raw | Dark |
| 68,39 | Cranberries. raw | Red |
| 5,11 | Cranberry bush berries. raw | Red |
| 157,78 | Currants. european black. raw | Dark |
| 109,62 | Currants. golden. raw | Dark |
| 75,02 | Currants. red. raw | Red |
| 85,69 | Eggplant. raw | Dark |
| 485,28 | Elderberries. raw | Dark |
| 31 | European cranberry | Red |

| | | |
|---|---|---|
| 407,6 | Evergreen huckleberry | Dark |
| 9,51 | Gooseberries. raw | Red |
| 72,1 | Grape [Black] | Red |
| 120,1 | Grapes. Concord. raw | Dark |
| 48,04 | Grapes. red. raw | Red |
| 169,17 | Half-highbush blueberry | Dark |
| 164,37 | Highbush blueberry, raw | Dark |
| 133,99 | Highbush blueberry. raw | Dark |
| 74 | Jostaberry | Dark |
| 27,88 | Jostaberry. raw | Dark |
| 40,15 | Lingonberries (cowberries). raw | Red |
| 45 | Lingonberry, raw | Red |
| 60,21 | Lingonberry. raw | Red |
| 149,17 | Lowbush blueberry, raw | Dark |
| 187,23 | Lowbush blueberry. raw | Dark |
| 88,52 | Maqui (Chilean wineberry). raw | Dark |
| 94,24 | Molucca raspberry. raw | Red |
| 24,77 | Muntries (emu apple. native cranberry. or munthar). raw | Red |
| 275,29 | Oval-leaf huckleberry | Dark |
| 752,68 | Peppers. tasmanian | Dark |
| 48,96 | Plum. Davidson | Red |
| 558,19 | Plum. Illawara. raw | Dark |
| 56,05 | Plums. black diamond. with peel. raw | Dark |
| 23,14 | Plums. purple. raw | Red |
| 138,71 | Rabbiteye blueberry | Dark |
| 686,79 | Raspberries. black | Dark |
| 48,63 | Raspberries. raw | Red |
| 16,5 | Red huckleberry | Red |
| 43,57 | Red raspberry, raw | Red |
| 72,49 | Red raspberry. raw | Red |
| 26,37 | Redcurrant, raw | Red |
| 38,58 | Redcurrant. raw | Red |
| 180,78 | Service (Saskatoon) berries | Dark |
| 358 | Skunk | Dark |
| 27,01 | Strawberries. raw | Red |
| 73 | Strawberry. raw | Red |

Wilcoxon rank sum test with continuity correction

data:  value by color
W = 1767, p-value = 1.08e-13
alternative hypothesis: true location shift is not equal to 0

| Color | N | Mean | Median | SD | IQR | Min | Max |
|---|---|---|---|---|---|---|---|
| Dark | 48 | 273.47167 | 170.880 | 248.39105 | 180.0800 | 27.88 | 1316.65 |
| Red | 38 | 38.67053 | 33.985 | 25.62553 | 27.1925 | 1.22 | 94.24 |
