## Supplementary material for "*Rubus armeniacus* genome sequence reveals the secrets of blackberry anthocyanin biosynthesis": Figure S1

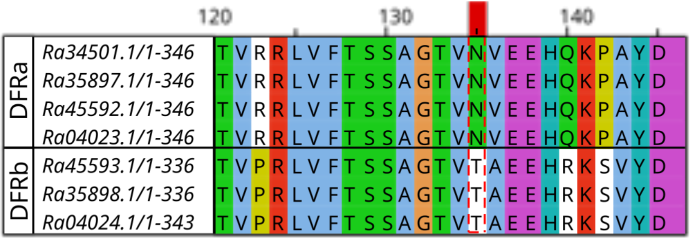


MAFFT alignment of DFRa and DFRb sequences against each other. The relevant amino achid for substrate specifity is framed with a red dotted line.
