## Supplementary material for "*Rubus armeniacus* genome sequence reveals the secrets of blackberry anthocyanin biosynthesis": Figure S2

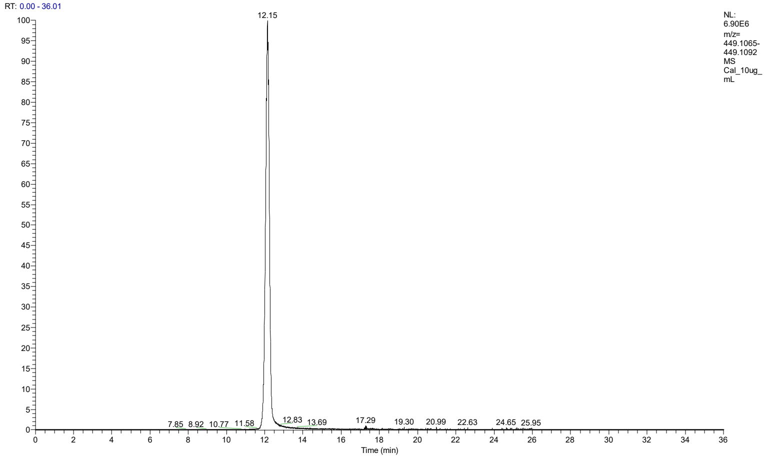


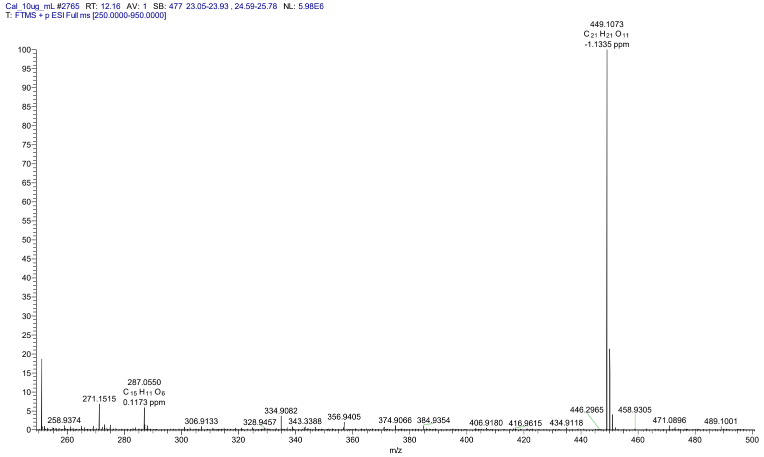


Reference: Cyanidin-3-glucoside 10 µg/mL
HPLC separation and HRAM MS
Mw: $C_{21}H_{21}O_{11}^{+}$ /// m/z 449.1084

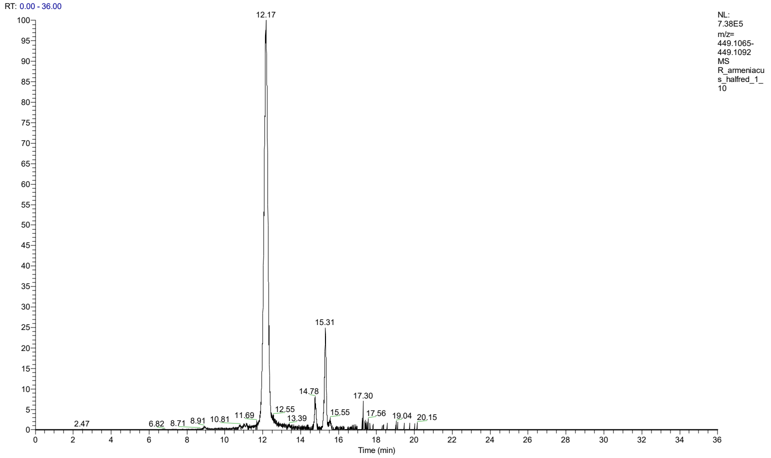


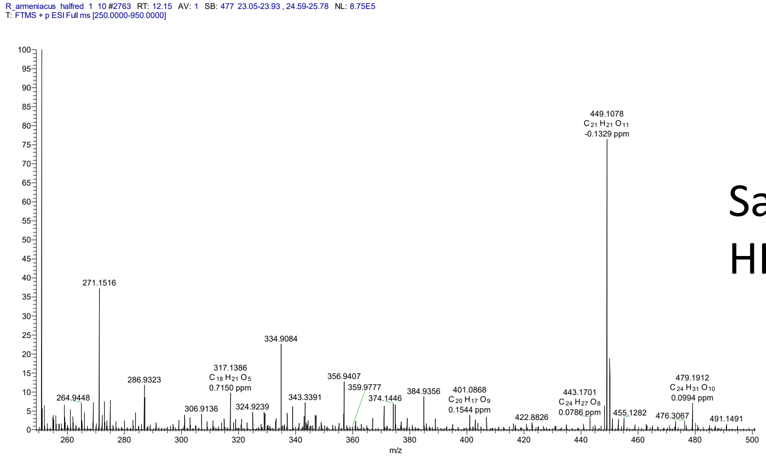


Sample: half-red blackberry extract diluted 1/10
HPLC separation and HRAM MS

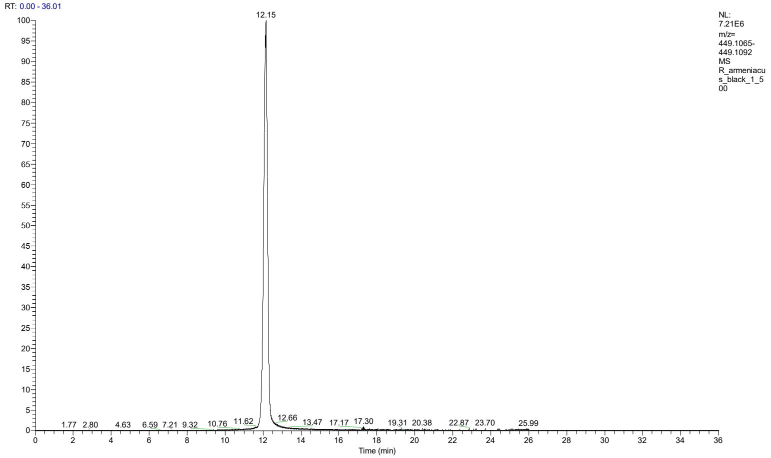


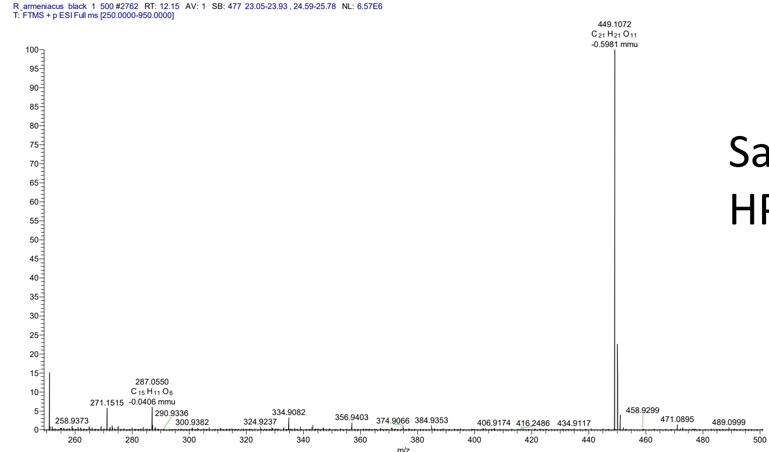


Sample: black blackberry extract diluted 1/500
HPLC separation and HRAM MS
