## Supplementary figures and images for "*Rubus armeniacus* genome sequence reveals the secrets of blackberry anthocyanin biosynthesis"

### Figure S3

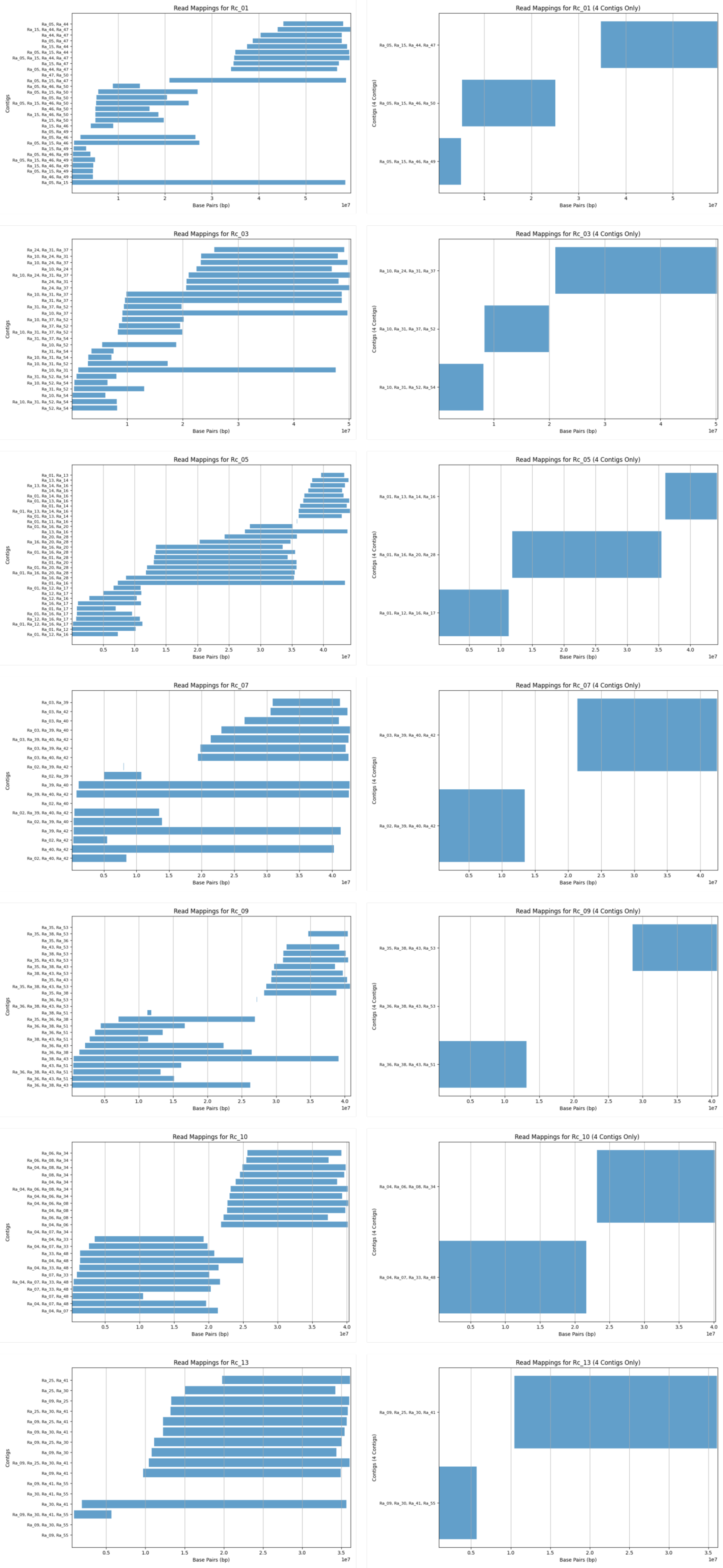
