## Supplementary material for "*Rubus armeniacus* genome sequence reveals the secrets of blackberry anthocyanin biosynthesis": Table S1

| Haplophase | contig | size in bp |
| --- | --- | --- |
| A | Ra_05 | 60730349 |
| A | Ra_10 | 51683244 |
| A | Ra_16 | 50081593 |
| A | Ra_39 | 45264764 |
| A | Ra_38 | 46949776 |
| A | Ra_04 | 41999628 |
| A | Ra_09 | 42464733 |
| B | Ra_15 | 56394652 |
| B | Ra_59 | 53147 |
| B | Ra_62 | 143644 |
| B | Ra_37 | 50729503 |
| B | Ra_12 | 11048478 |
| B | Ra_18 | 322204 |
| B | Ra_61 | 2000486 |
| B | Ra_01 | 37136298 |
| B | Ra_40 | 41259553 |
| B | Ra_57 | 50738 |
| B | Ra_43 | 35485994 |
| B | Ra_23 | 71565 |
| B | Ra_65 | 281115 |
| B | Ra_07 | 28852864 |
| B | Ra_08 | 16241447 |
| B | Ra_41 | 41254849 |
| C | Ra_46 | 24540304 |
| C | Ra_44 | 23989297 |
| C | Ra_52 | 20579474 |
| C | Ra_22 | 92204 |
| C | Ra_64 | 637254 |
| C | Ra_24 | 35456637 |
| C | Ra_17 | 11257227 |
| C | Ra_26 | 153851 |
| C | Ra_27 | 162883 |
| C | Ra_28 | 25996246 |
| C | Ra_11 | 105176 |
| C | Ra_14 | 7713660 |
| C | Ra_02 | 12809766 |
| C | Ra_63 | 988063 |
| C | Ra_56 | 55268 |
| C | Ra_03 | 24086050 |
| C | Ra_36 | 30768823 |
| C | Ra_53 | 19280844 |
| C | Ra_29 | 193281 |
| C | Ra_48 | 19740449 |
| C | Ra_34 | 18350832 |
| C | Ra_55 | 5809857 |
| C | Ra_25 | 31145472 |
| D | Ra_49 | 4571763 |
| D | Ra_60 | 78289 |
| D | Ra_50 | 17111329 |

| | | |
|---|---|---|
| D | Ra_47 | 23761129 |
| D | Ra_31 | 43951974 |
| D | Ra_20 | 25760182 |
| D | Ra_13 | 7523004 |
| D | Ra_58 | 50726 |
| D | Ra_42 | 33795919 |
| D | Ra_51 | 18547055 |
| D | Ra_45 | 609109 |
| D | Ra_35 | 14673032 |
| D | Ra_32 | 55540 |
| D | Ra_33 | 28039320 |
| D | Ra_06 | 20661203 |
| D | Ra_21 | 266096 |
| D | Ra_19 | 251885 |
| D | Ra_30 | 30642381 |
| D | Ra_54 | 8577700 |
