## Supplementary material for "*Rubus armeniacus* genome sequence reveals the secrets of blackberry anthocyanin biosynthesis": Table S3

| ID | Gene | Similarity | ConservedResidues | ConservedRegions |
|---|---|---|---|---|
| Ra50464.1 | C4H_1 | 81.52972833254675 | 100.0 | 100.0 |
| Ra63304.1 | C4H_2 | 81.47445671127444 | 100.0 | 100.0 |
| Ra28206.1 | C4H_3 | 81.47015186604867 | 100.0 | 100.0 |
| Ra71901.1 | C4H_4 | 81.4105176075289 | 100.0 | 100.0 |
| Ra65568.1 | F3-H_1 | 70.80225422789373 | 100.0 | 87.51157407407408 |
| Ra22443.1 | F3-H_2 | 70.73008855361311 | 100.0 | 87.51157407407408 |
| Ra18381.1 | F3-H_3 | 70.51008694033011 | 100.0 | 87.51157407407408 |
| Ra09034.1 | F3-H_4 | 70.40057483141187 | 100.0 | 87.51157407407408 |
| Ra38240.1 | F3-5-H_1 | 37.39717807507978 | 100.0 | 54.47475608600928 |
| Ra14784.1 | F3-5-H_2 | 37.27778219497314 | 100.0 | 54.47475608600928 |
| Ra100611.1 | F3-5-H_3 | 37.19438900179078 | 100.0 | 54.47475608600928 |
| Ra108198.1 | F3-5-H_4 | 37.17647963267918 | 100.0 | 54.47475608600928 |
| Ra59989.1 | FNS2_1 | 49.09723025445479 | 100.0 | 85.14285714285714 |
| Ra31412.1 | FNS2_2 | 48.73513585572547 | 100.0 | 85.14285714285714 |
| Ra65568.1 | FNS2_3 | 36.22442475337656 | 100.0 | 73.28571428571429 |
| Ra22443.1 | FNS2_4 | 36.10551115249361 | 100.0 | 73.28571428571429 |
| Ra18381.1 | FNS2_5 | 35.89327336264658 | 100.0 | 73.28571428571429 |
| Ra09034.1 | FNS2_6 | 35.87471161836435 | 100.0 | 73.28571428571429 |
| Ra34501.1 | DFR_1 | 68.23497778942087 | 100.0 | 80.51839464882943 |
| Ra35897.1 | DFR_2 | 68.23497778942087 | 100.0 | 80.51839464882943 |
| Ra45592.1 | DFR_3 | 67.74876087688486 | 100.0 | 80.51839464882943 |
| Ra04023.1 | DFR_4 | 68.02861666113368 | 100.0 | 78.34448160535118 |
| Ra45593.1 | DFR_5 | 65.78615195057866 | 100.0 | 72.5752508361204 |
| Ra35898.1 | DFR_6 | 65.38503168370674 | 100.0 | 72.5752508361204 |
| Ra04024.1 | DFR_7 | 65.45460450551737 | 100.0 | 70.40133779264214 |
| Ra59429.1 | ANR_1 | 76.40555265830682 | 100.0 | 67.53246753246754 |
| Ra31968.1 | ANR_2 | 76.22370417690026 | 100.0 | 67.53246753246754 |
| Ra116680.1 | ANR_3 | 76.22370417690026 | 100.0 | 67.53246753246754 |
| Ra111735.1 | ANR_4 | 76.10979071565062 | 100.0 | 67.53246753246754 |
| Ra93173.1 | CHS_1 | 86.95331415339207 | 100.0 | 94.11764705882354 |
| Ra04232.1 | CHS_2 | 86.89276528468585 | 100.0 | 94.11764705882354 |
| Ra86200.1 | CHS_3 | 86.80988364035464 | 100.0 | 94.11764705882354 |
| Ra78742.1 | CHS_4 | 86.72795994484062 | 100.0 | 94.11764705882354 |
| Ra04231.1 | CHS_5 | 86.54615783600693 | 100.0 | 94.11764705882354 |
| Ra86201.1 | CHS_6 | 86.51479681991702 | 100.0 | 94.11764705882354 |
| Ra93174.1 | CHS_7 | 86.45201253648656 | 100.0 | 94.11764705882354 |
| Ra78741.1 | CHS_8 | 86.3811990385707 | 100.0 | 94.11764705882354 |
| Ra25478.1 | F3H_1 | 83.04964552466203 | 100.0 | 95.28769841269842 |
| Ra51951.1 | F3H_2 | 83.04964552466203 | 100.0 | 95.28769841269842 |
| Ra56921.1 | F3H_3 | 83.04964552466203 | 100.0 | 95.28769841269842 |
| Ra88419.1 | F3H_4 | 83.04964552466203 | 100.0 | 95.28769841269842 |
| Ra33387.1 | FLS_1 | 67.98986057213955 | 100.0 | 0.0 |
| Ra44441.1 | FLS_2 | 67.90235019061346 | 100.0 | 0.0 |
| Ra34743.1 | FLS_3 | 67.87630523663887 | 100.0 | 0.0 |
| Ra02881.1 | FLS_4 | 67.70051287516515 | 100.0 | 0.0 |
| Ra23521.1 | ANS_1 | 74.36632291050105 | 100.0 | 0.0 |
| Ra10125.1 | ANS_2 | 73.94873479200959 | 100.0 | 0.0 |
| Ra19492.1 | ANS_3 | 73.94873479200959 | 100.0 | 0.0 |
| Ra66679.1 | ANS_4 | 73.92437238588774 | 100.0 | 0.0 |
| Ra68780.1 | CHI1_1 | 68.44480125708888 | 100.0 | 0.0 |
| Ra77528.1 | CHI1_2 | 68.44464195647315 | 100.0 | 0.0 |
| Ra94348.1 | CHI1_3 | 68.44464195647315 | 100.0 | 0.0 |
| Ra115050.1 | CHI1_4 | 68.36032650069602 | 100.0 | 0.0 |

| | | | | |
|---|---|---|---|---|
| Ra86843.1 | TT19_1 | 59.88532559169914 | 100.0 | 0.0 |
| Ra58529.1 | TT19_2 | 59.72430341203695 | 100.0 | 0.0 |
| Ra27175.1 | TT19_3 | 59.697580592485195 | 100.0 | 0.0 |
| Ra53627.1 | TT19_4 | 59.697580592485195 | 100.0 | 0.0 |
| Ra108874.1 | PAL_1 | 82.39644316880305 | 100.0 | 100.0 |
| Ra101362.1 | PAL_2 | 82.36868026572212 | 100.0 | 100.0 |
| Ra37530.1 | PAL_3 | 82.3060840987212 | 100.0 | 100.0 |
| Ra14120.1 | PAL_4 | 82.22295849533738 | 100.0 | 100.0 |
| Ra114897.1 | PAL_5 | 79.37443718422945 | 100.0 | 100.0 |
| Ra68917.1 | PAL_6 | 79.24481410105619 | 100.0 | 100.0 |
| Ra77375.1 | PAL_7 | 79.24481410105619 | 100.0 | 100.0 |
| Ra94493.1 | PAL_8 | 79.20576113270872 | 100.0 | 100.0 |
| Ra06378.1 | LAR_1 | 64.17883530113177 | 100.0 | 100.0 |
| Ra83291.1 | LAR_2 | 64.1591865514883 | 100.0 | 100.0 |
| Ra81676.1 | LAR_3 | 63.94608988510843 | 100.0 | 100.0 |
| Ra90382.1 | LAR_4 | 63.82561511652732 | 100.0 | 100.0 |
| Ra05133.1 | 4CL_1 | 68.84085797870556 | 100.0 | 97.72727272727272 |
| Ra85394.1 | 4CL_2 | 68.77649321946998 | 100.0 | 97.72727272727272 |
| Ra79562.1 | 4CL_3 | 68.70899152879886 | 100.0 | 97.72727272727272 |
| Ra92424.1 | 4CL_4 | 68.34467731888206 | 100.0 | 97.72727272727272 |
| Ra97346.1 | 4CL_5 | 68.7197250953251 | 100.0 | 90.9090909090909 |
| Ra38926.1 | 4CL_6 | 68.65774544311239 | 100.0 | 90.9090909090909 |
| Ra105309.1 | 4CL_7 | 68.64544963627243 | 100.0 | 90.9090909090909 |
| Ra15428.1 | 4CL_8 | 68.61677096454416 | 100.0 | 90.9090909090909 |
