## Supplementary material for "*Rubus armeniacus* genome sequence reveals the secrets of blackberry anthocyanin biosynthesis": Table S4

Comparison of available genome sequences for the genus *Rubus* with an assembly level of scaffold or higher. Additionally, the statistics for the genome sequence of *Rubus armeniacus* generated by this study are included. NCBI reference assemblies are marked as * behind the species name. All BUSCO scores are calculated based on the dataset rosacea_odb12.

| **Assembly Accession** | **Species** | **Assembly Level** | **Number of Scaffolds** | **LAI** | **BUSCO** | **Reference** |
|---|---|---|---|---|---|---|
| GCA_051990025.1 | *Rubus alceifolius* * | complete | 14 | 13.58 | C:99.0%<br>S:10.7%, D:88.3% | DOI:10.1101/2025.04.09.646780 |
| GCA_964235055.1 | *Rubus caesius* * | chromosome | 428 | 8.48 | C:99.1%<br>S:3.4%, D:95.6% | Darwin Tree of Life |
| GCA_040183295.1 | *Rubus argutus Hillquist* * | chromosome | 350 | 8.67 | C:98.1%<br>S:89.2%, D:8.9% | DOI:10.1093/g3journal/jkac289 |
| GCA_030142095.1 | *Rubus idaeus Malling Jewel* | chromosome | 11 | 5.14 | C:98.2%<br>S:95.4%, D:2.9% | DOI: 10.1371/journal.pone.0285756 |
| GCA_965153425.1 | *Rubus idaeus* | chromosome | 61 | 8.14 | C:98.8%<br>S:96.7%, D:2.0% | Darwin Tree of Life |
| GCA_030142025.1 | *Rubus idaeus Autumn Bliss* | chromosome | 17 | 4.83 | C:98.3%<br>S:90.7%, D:7.6% | DOI: 10.1371/journal.pone.0285756 |
| GCA_965153365.1 | *Rubus idaeus* | scaffold | 164 | 7.21 | C:98.8%<br>S:96.7%, D:2.2% | Darwin Tree of Life |
| GCA_029955405.1 | *Rubus parviflorus* * | scaffold | 235 | 5.50 | C:97.5%<br>S:94.5%, D:3.0% | - |

| | | | | | | |
|---|---|---|---|---|---|---|
| GCA_964235075.1 | *Rubus caesius* | scaffold | 295 | 7.54 | C:99.1% S:3.7%, D:95.4% | Darwin Tree of Life |
| GCA_047496405.1 | *Rubus idaeus Varnes* * | scaffold | 13 | 5.47 | C:98.7% S:96.4%, D:2.3% | - |
| GCA_029955375.1 | *Rubus parviflorus* | scaffold | 149 | 9.87 | C:98.2% S:94.8%, D:3.4% | - |
| GCA_040496155.1 | *Rubus hochstetterorum* * | scaffold | 1114283 | n.a | C:17.7% S:17.1%, D:0.6% | - |
| Whole assembly | *Rubus armeniacus* | contig | 65 | 19.54 | C:99.1% S:0.2%, D:98.9% | this study |
| hapA | *Rubus armeniacus* | complete | 7 | 24.72 | C:98.8% S:97.1%, D:1.8% | this study |
